# Spatiotemporal organization of human sensorimotor beta burst activity

**DOI:** 10.1101/2022.05.19.492617

**Authors:** Catharina Zich, Andrew J Quinn, James J Bonaiuto, George O’Neill, Lydia C Mardell, Nick S Ward, Sven Bestmann

## Abstract

Beta oscillations in human sensorimotor cortex are hallmark signatures of healthy and pathological movement. In single trials, beta oscillations include bursts of intermittent, transient periods of high-power activity. These burst events have been linked to a range of sensory and motor processes, but their precise spatial, spectral, and temporal structure remains unclear. Specifically, a role for beta burst activity in information coding and communication suggests spatiotemporal patterns, or travelling wave activity, along specific anatomical gradients. We here show in human magnetoencephalography recordings that burst activity in sensorimotor cortex occurs in planar spatiotemporal wave-like patterns that dominate along two axes either parallel or perpendicular to the central sulcus. Moreover, we find that the two propagation directions are characterised by distinct anatomical and physiological features. Finally, our results suggest that sensorimotor beta bursts occurring before and after a movement share the same generator but can be distinguished by their anatomical, spectral and spatiotemporal characteristics, indicating distinct functional roles.

## Introduction

Neural activity at the rate of 13-30Hz constitute one of the most prominent electrophysiological signatures in the sensorimotor system (Baker, 2007; Brown, 2007). This sensorimotor beta activity is traditionally seen to reflect oscillations: sustained rhythmic synchronous spiking activity within neural populations. However, a substantial proportion of sensorimotor beta activity occurs in bursts of intermittent, transient periods of synchronous spiking activity (Jones, 2016) which relate to both motor, perceptual and sensory function (Enz et al., 2021; Feingold et al., 2015; Heideman et al., 2020; Sherman et al., 2016; Shin et al., 2017; Sporn et al., 2020; Tinkhauser, Pogosyan, Little, et al., 2017; Wessel, 2020; Zich et al., 2018) and pathophysiological movement (Cagnan et al., 2019; Deffains et al., 2018; Tinkhauser, Pogosyan, Little, et al., 2017; Tinkhauser, Pogosyan, Tan, et al., 2017), but their functional role remains unclear.

Sensorimotor beta burst activity is commonly considered as zero-lagged (or standing wave) activity which is generated by the summation of synchronized layer-specific inputs within cortical columns that result in a cumulative dipole with a stereotypical wavelet shape in the time domain (Bonaiuto et al., 2021; Law et al., 2022; Neymotin et al., 2020). These time-periods of synchronous activity which generate standing wave activity are thought to convey little information encoding (Brittain & Brown, 2014; Carhart-Harris, 2018; Carhart-Harris et al., 2014). This view sides with the proposed akinetic role of high sensorimotor beta states (Gilbertson et al., 2005; Joundi et al., 2012; Khanna & Carmena, 2017; Pogosyan et al., 2009). However, burst activity may have heterogenous and mechanistically distinct components which can be characterised by their distinct spatial, temporal, and spectral structure (Law et al., 2022; Zich et al., 2020) that, in addition to zero-lagged activity, contains spatiotemporal gradients, or travelling wave, components.

In animals, for example, a high proportion of sensorimotor beta activity occurs as travelling waves (Rubino et al., 2006; Rule et al., 2018), in addition to highly synchronous standing waves. In travelling waves, the relative timing of fluctuations of synchronous spiking activity is not precisely zero-lagged but adopts a phase offset and moves across space. Propagation of neural activity constitutes one mechanism for cortical information transfer and traveling waves have been described over spatial scales that range from the mesoscopic (single cortical areas and millimetres of cortex) to the macroscopic (global patterns of activity over several centimetres) and extend over temporal scales from tens to hundreds of milliseconds (Alexander et al., 2019; Davis et al., 2021; Heitmann et al., 2017; Muller et al., 2018; Roberts et al., 2019; Rule et al., 2018).

Characterising traveling wave components within sensorimotor beta burst activity is of relevance as it would provide insights into the putative underlying mechanisms and functional roles of sensorimotor beta activity. For instance, in general terms, spatiotemporal propagation of high amplitude beta may support information transfer across space and may reflect the spatiotemporal patterns of sequential activation required for movement initiation (Best et al., 2016; Rubino et al., 2006). At the macro-scale level, the specific propagation properties, such as propagation direction and speed, may provide further constraints for the putative functional role of burst activity in organizing behaviour across different brain regions (Ding & Ermentrout, 2021), including the modulation of neural sensitivity (Davis et al., 2020) or the sequencing of muscle representations in motor cortex (Muller et al., 2018; Riehle et al., 2013; Takahashi et al., 2015). In humans, the precise properties of beta bursts, and whether their high amplitude activity comprise distinct spatiotemporal gradients remains unclear.

To address this, we here employed high signal-to-noise (SNR) magnetoencephalography (MEG) in healthy human subjects during simple visually-cued motor behaviour. We show that beta burst activity in sensorimotor cortex occurs in planar spatiotemporal wave-like patterns that dominate along two anatomical axes. Crucially, our results show structure beyond the inherent limitations of source reconstruction such as volume conduction or the spatial pattern of beamformer weights. Moreover, we find that the two propagation directions are characterised by distinct anatomical and physiological features. Finally, our results suggest that sensorimotor beta bursts occurring before and after a movement share the same generator but can be distinguished by their anatomical, spectral and spatiotemporal characteristics, indicating distinct functional roles.

## Results

### Temporal, spectral and spatial burst characteristics

Participants completed three blocks per recording session, and 1-5 sessions on different days. We analysed 123-611 trials per participant (*M* = 438.5, *SD* = 151.0 across individuals) in which correct key presses were made with either the right index or middle finger, in response to congruent imperative stimuli and high coherence visual cues (Bonaiuto et al., 2018; Little et al., 2019). We focussed on these trial-types to delineate the multi-dimensional (temporal, spectral, spatial) properties of sensorimotor beta burst activity (**Fig. 1a,b**; (Zich et al., 2020)). Bursts were identified over a 4 second time window (−2 to 2s relative to the button press), in the beta frequency range (13 to 30Hz) and a region-of-interest (ROI) spanning the primary motor cortex (M1) and adjacent areas of the primary sensory cortex and premotor cortex using session-specific amplitude thresholding ((Little et al., 2019); **Supplemental Fig. 1**; **Supplemental Fig. 2**) and 5D clustering.

**Fig. 1:**
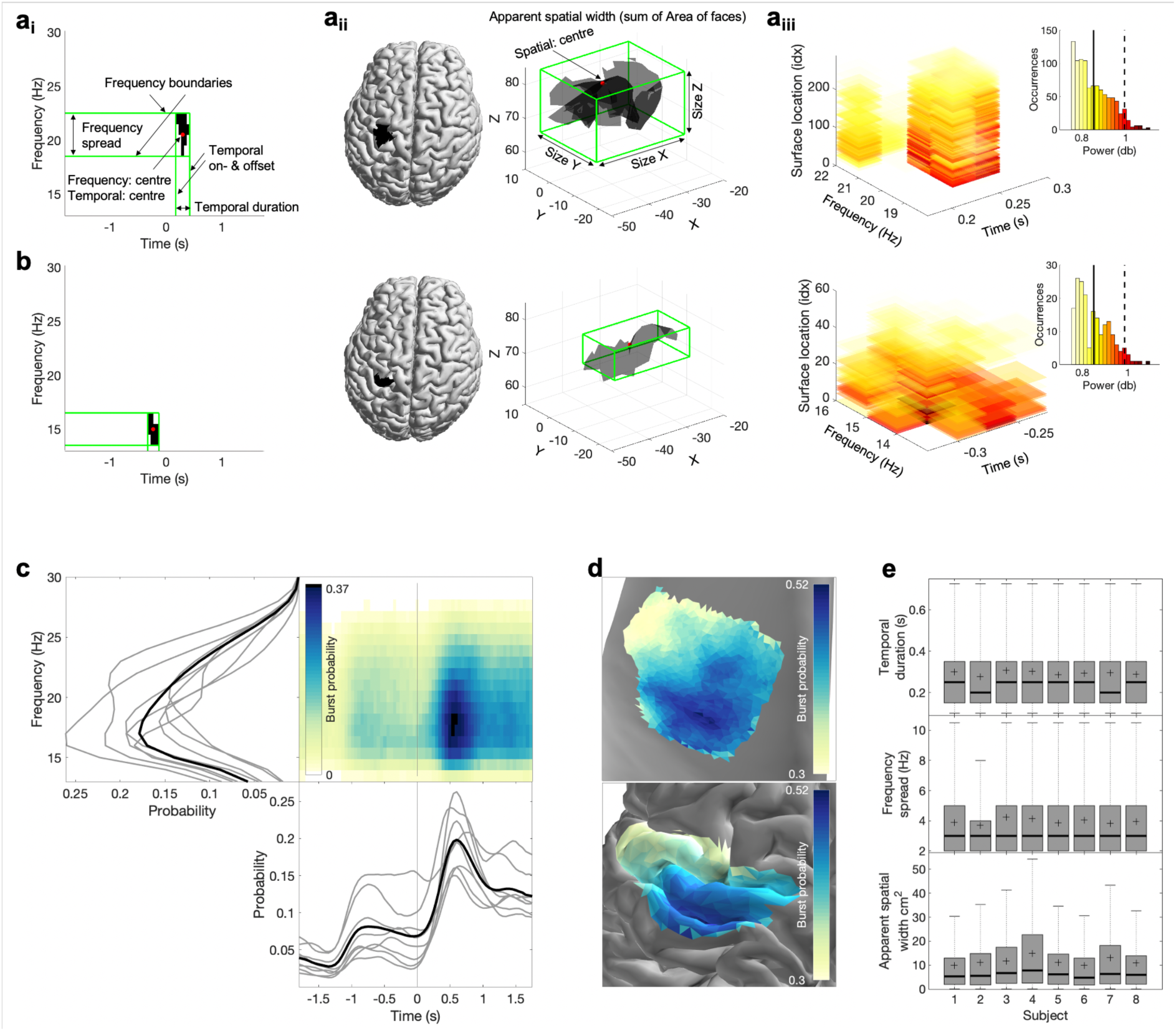
Spectral, temporal, and spatial beta burst characteristics. (a) Burst characteristics for a single example burst. (a_i_) Temporal and spectral burst characteristics. (a_ii_) Spatial burst characteristics. (a_iii_) Burst amplitude. Shown is the amplitude for each temporal, spectral and spatial location of that burst as well as the histogram across all three signal domains. Often the mean (straight line) or the 95 percentile (dashed line) is used as burst amplitude. (b) Same as (a) for a different burst. (c) Burst probability as a function of time and frequency across all bursts of all subjects (see **Supplemental Fig. 3** for individual subjects). Burst probability as a function of time (bottom) and frequency (left) is shown for each subject separately (grey lines) and across subjects (black line). (d) Burst probability as a function of space across all bursts of all subjects on the inflated surface (top) and original surface (bottom). To visualise burst probability as a function of space across subjects, individual subject maps were spatially normalised, projected onto a single surface, and then averaged across subjects. **Supplemental Fig. 3** depicts the burst probability for each subject in native space. (e) Burst temporal duration (top), frequency spread (middle) and apparent spatial width (bottom) for each subject as boxplot.

In the temporal domain, we observed the expected increase in burst probability post-vs pre-movement (**Fig. 1c**). Burst duration was consistent across subjects (*M* = 238ms, *SD* = 23ms across individuals, temporal resolution 50ms, **Fig. 1e**). Spectrally, while beta bursts occurred throughout the beta frequency range, most bursts were identified in the lower beta frequency range (**Fig. 1c**), with a consistent frequency spread across subjects (*M* = 3Hz, *SD* = 0Hz across individuals, frequency resolution 1Hz, **Fig. 1e**). To examine burst probability as a function of space across subjects, individual subject maps were spatially normalised, projected onto a single surface, and then averaged across subjects. Topographically, bursts were most likely to occur in M1 (**Fig. 1d**, see **Supplemental Fig. 3** for individual subject maps) and spanned, on average, 10% of the ROI’s surface area (*M* = 6cm^2^*; SD* = 0.9cm^2^ across individuals).

We performed a range of control analyses to examine whether our results can be explained by trivial properties of the beamformer itself. Firstly, we sought to assess whether differences in the bursts’ apparent spatial width could be explained by differences in SNR across and/or within sessions rather than differences in the spatial distribution of cortical activity. We reasoned that if differences in SNR across sessions would explain bursts’ apparent spatial width, then burst amplitude and burst apparent spatial width should be negatively correlated (for a schematic illustration see **Supplemental Fig. 4ai**). The absence of significant correlations between burst amplitude and burst apparent spatial width, both across sessions within subjects and also across sessions and individuals (**Supplemental Fig. 4aii**), suggests that the apparent spatial width of bursts is not solely explained by differences in SNR across sessions, and across individuals.

Further we reasoned that if the apparent spatial width is driven by differences in SNR across bursts within a session, then a positive relationship between burst amplitude and burst apparent spatial width within sessions should be present, and there should be no systematic phase differences across different spatial locations within each burst (for a schematic illustration see **Supplemental Fig. 4b**). While burst amplitude and burst apparent spatial width are positively correlated within sessions (Pearson’s *r*: *M* = 0.749, *SD* = 0.056 across sessions, all *p’s* < 0.001), we consistently observed diverse phase lags across space within these bursts (see results section: **Sensorimotor burst activity propagates along two axes**), which are unlikely to arise simply from amplitude scaling of a single source.

Together, these control analyses suggest that differences in bursts’ apparent spatial width is not merely due to differences in SNR across and/or within sessions, but for the most part due to differences in the spatial distribution of cortical activity.

### Sensorimotor beta burst activity is propagating

The precise decomposition of beta bursts into their spectral, spatial, and temporal signal domains allowed us to next assess any spatiotemporal gradients within sensorimotor beta bursts. For each burst, we identified the dominant propagation direction and propagation speed (**Supplemental Fig. 1**). Propagation direction and speed were estimated from critical points in the oscillatory cycle (**Fig. 2a**) and then averaged across critical points within one burst. The propagation direction at each critical point was estimated from the relative latency (**Fig 2bi**). Next, the propagation direction was estimated, using linear regression (Balasubramanian et al., 2020), whereby the relative latency at the surface location was predicted from the coordinates of the surface location of the inflated surface. We excluded complex spatiotemporal patterns such as random or circular patterns (**Supplemental Fig. 5b,c**; *M* = 8.25%, *SD* = 0.88% across individuals; (Denker et al., 2018; Rule et al., 2018)), to focus on bursts with a dominant planar propagation orientation (**Supplemental Fig. 5a**; *M* = 79.6%, *SD* = 2.4% across individuals; (Balasubramanian et al., 2020; Rubino et al., 2006; Takahashi et al., 2011)).

**Fig. 2:**
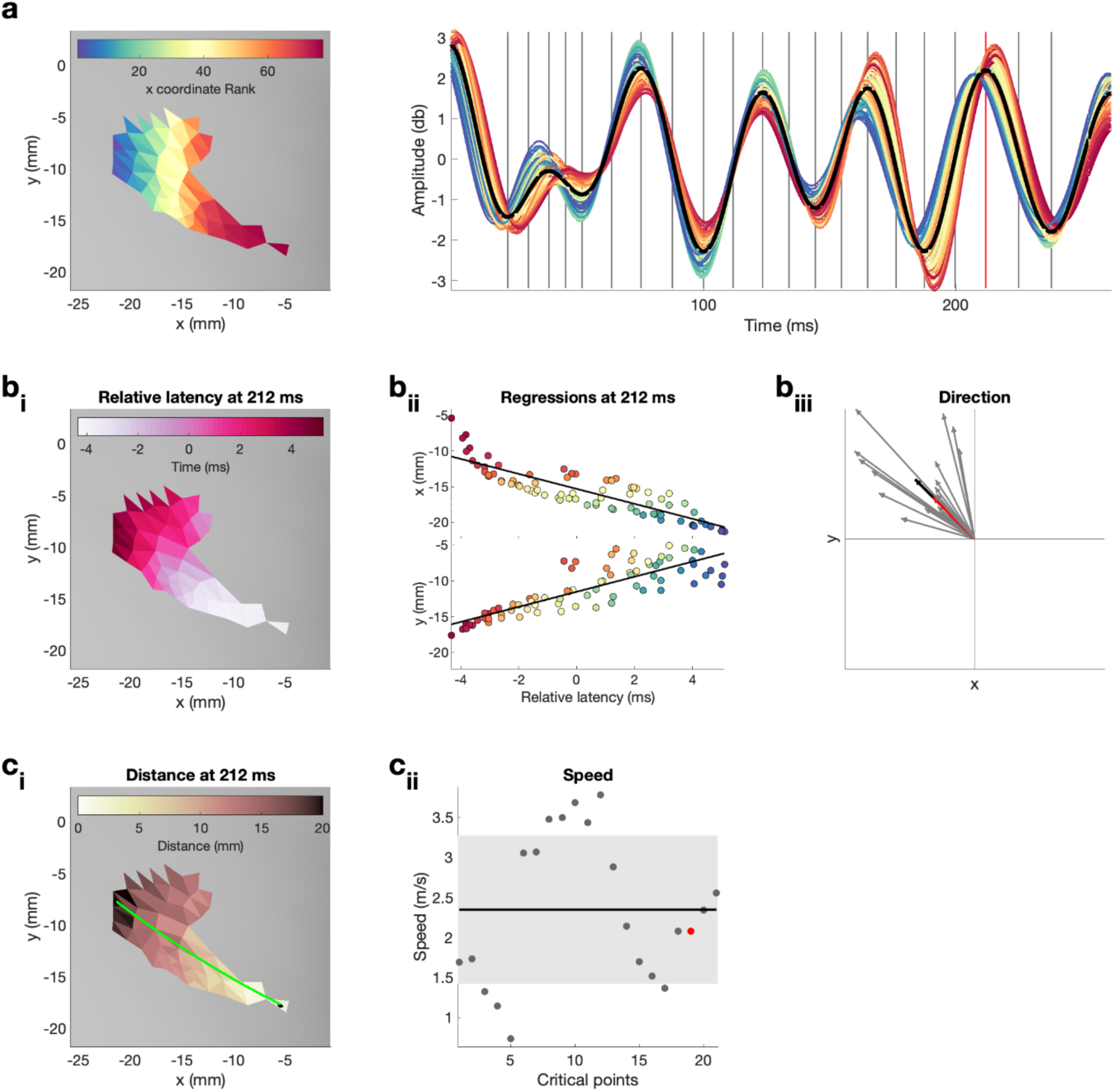
Quantification of propagation direction and propagation speed on one exemplar burst. For a dynamic version, i.e., updated for each critical point, see Supplemental Video 1. (a) Left: Single burst on inflated surface. Spatial locations are colour-coded by their x coordinate rank. Right: Neural activity in the beta range (13-30 Hz) from each surface location for the temporal duration of the burst. Vertical lines indicate critical points (four critical points per oscillatory cycle, i.e., peak and trough as well as peak-trough and trough-peak midpoint) at which propagation direction and propagation speed were estimated. Red vertical line indicates the control point at 212ms, shown in (b_i,ii_) and (c_i,ii_), and highlighted in (b_iii_) and (c_iii_). (b_i_) Relative latencies of the critical point at 212ms as a function of space illustrated on inflated surface. (b_ii_) Simple linear regressions between latency at surface location and x (top) as well as y (bottom) coordinates of the surface location for the critical point at 212ms. Colour refers to the x coordinate rank as illustrated in (a). (b_iii_) Propagation direction obtained from regression coefficients for each critical point (grey), the critical point at 212ms (red) and the average across all critical points (black, i.e., propagation direction of burst). (c_i_) Distance, i.e., exact geodesic distance, from the surface location with the smallest relative latency to each surface location on the inflated surface for the critical point at 212ms. Green line indicated the path, i.e., distance, from the surface location with the smallest to the surface location with the largest relative latency. (c_ii_) Propagation speed for each critical point (grey), the critical point at 212ms (red). The standard deviation across critical points is indicated by the grey area and the average across all critical points (i.e., propagation speed of burst) is indicated by the black horizontal line.

To test whether the planar spatiotemporal structure of bursts is significant we compared the propagation properties detected in real burst activity to those of surrogate data for a subset of 100 randomly selected bursts. For each burst, 1000 phase-randomised surrogates (cf. Hurtado et al., 2004) were created and the propagation properties of the real data were compared to their distribution from 1000 surrogates. Real sensorimotor beta burst activity exhibited significantly stronger planar spatiotemporal structure than spectrally matched surrogate data (all 100 bursts *p*<0.01).

### Accuracy of the propagation direction estimation in simulated and surface meshes

Before assessing the propagation direction of sensorimotor beta burst activity, we evaluated the accuracy of the propagation direction estimation. To this end, we created 360 noise free high-resolution gradients spanning 1deg-360deg (in steps of 1deg; subset shown in **Fig. 3a**). The propagation direction of these gradients was then estimated from three different 2D surface mesh types (square mesh, circular mesh, random mesh; **Fig. 3b-d**) and three spatial sampling rates (N/2, N, Nx2, whereby N approximates the spatial sampling of the real mesh). By comparing the true and estimated propagation direction we found that under noise free conditions, the propagation direction can be estimated accurately from regular meshes (**Fig. 3b,c**), whereas the mean error is roughly twice as large for random meshes (**Fig. 3d**).

**Fig. 3:**
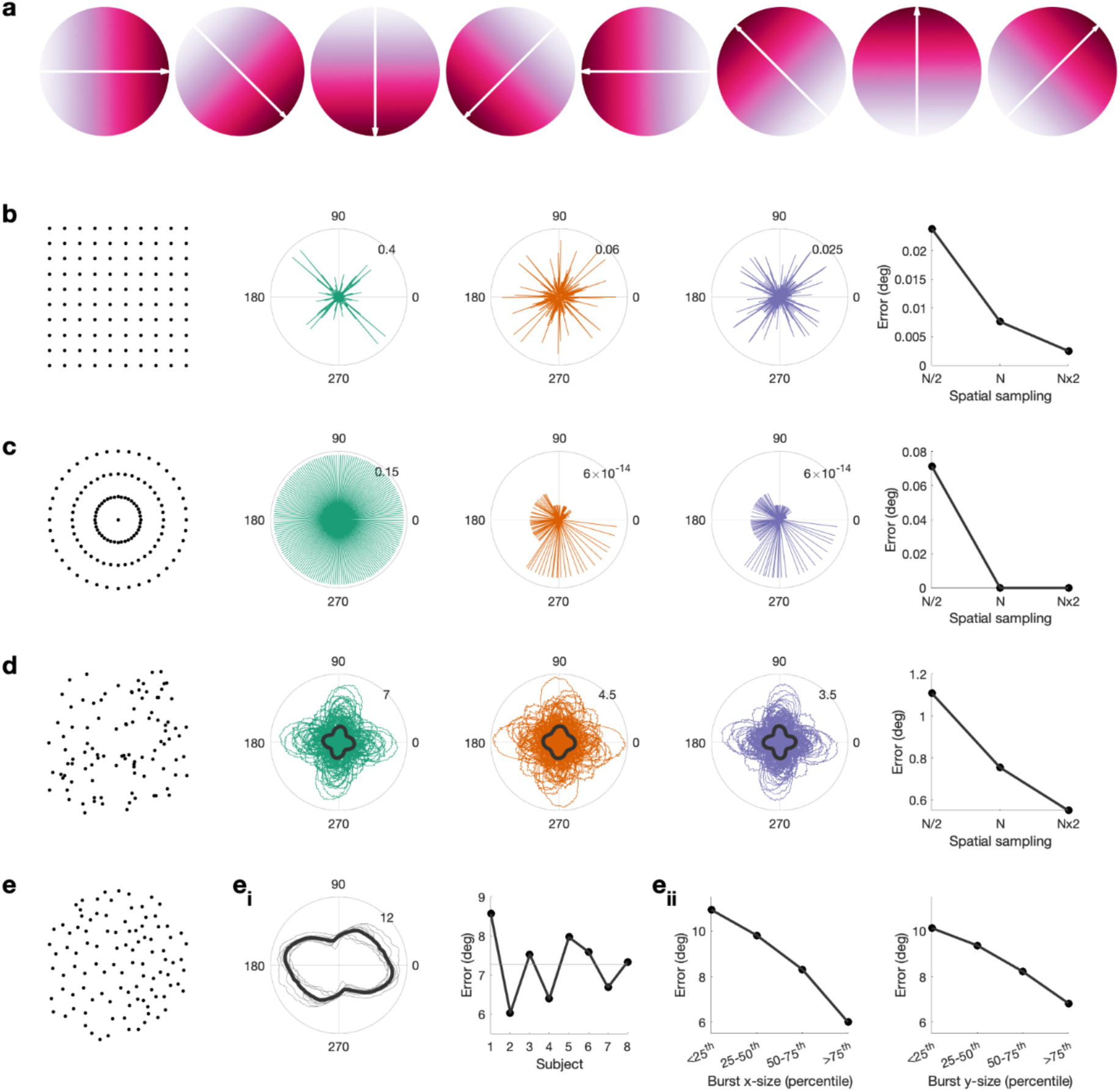
Propagation direction can be estimated accurately from cortical meshes. (a) Simulated gradient at 0, 45, 90, 135, 180, 225 and 270 degrees. (b) Error between simulated gradient and estimated gradient on a square mesh. We test three different spatial sampling rates, N/2 (green), N (red) and Nx2 (blue), whereby the spatial sampling of N is roughly equivalent to the spatial sampling of the inflated surface. Error is calculated for 1-360deg in steps of 1deg. The median error per spatial sampling is shown in the right, i.e., higher spatial sampling results in a lower error. (c) As (b), but for a circular mesh. (d) Error between simulated gradient and estimated gradient on a random mesh. 100 random meshes were generated. The error for each iteration is shown as well as the mean across iterations (black line). (e) Error between simulated gradient and estimated gradient on the inflated surface. The error was calculated for each burst. (ei) The mean across bursts is shown for each subject (grey lines) and across subjects (black line). For each subject the mean error across all angles and bursts is shown, i.e., error is comparable across subjects. (eii) Error as a function of burst size along the x-axis (left) and y-axis (right), i.e., the error is lower in bigger burst.

This is relevant because the surface mesh obtained from brain imaging data is irregular. When evaluating the accuracy of the propagation direction estimates using the real mesh (**Fig. 3e**) and the real spatial burst properties, we found an average error of 7deg between true and estimated propagation direction, with little variability across individuals (*SD* = 0.5deg across subjects) and angles (*SD* = 1.5deg across angles; **Fig. 3e_i_**). Across individuals the error was smallest for gradient directions around 104/284deg and largest around 170/350deg (**Fig. 3e_i_**). Further, for bursts with a larger apparent spatial width (i.e., containing more spatial samples), the estimated error is lower (**Fig. 3e_ii_**).

Overall, these results suggest that propagation directions can be estimated with sufficient accuracy from higher SNR MEG recordings over a relatively small cortical patch, as here.

### Sensorimotor burst activity propagates along two axes

Having established that the spatial sampling of the cortical mesh is sufficient to detect propagation in simulated gradients, we analysed the propagation properties of the sensorimotor beta burst activity. We observed that neural activity within beta bursts propagates along two dominant axes, which were approximately 90deg apart (**Fig. 4a**): one anterior-posterior (a-p) axis traversing the central sulcus in approximately perpendicular fashion, and one medial-lateral (m-l) axis running approximately parallel to (i.e., along) the central sulcus. The propagation distributions along these axes were well described by a mixture of four von Mises functions with means of 66deg and 248deg for the a-p axis, and means of 142deg and 324deg for the m-l axis, indicating that the surface mesh imposes structure. Note, however, that these axes do not align with the directions showing the smallest or the largest error when estimating the direction from noise-free gradients on the same surface mesh and spatial burst properties, indicating that the mesh properties do not drive the observed propagation direction.

**Fig. 4:**
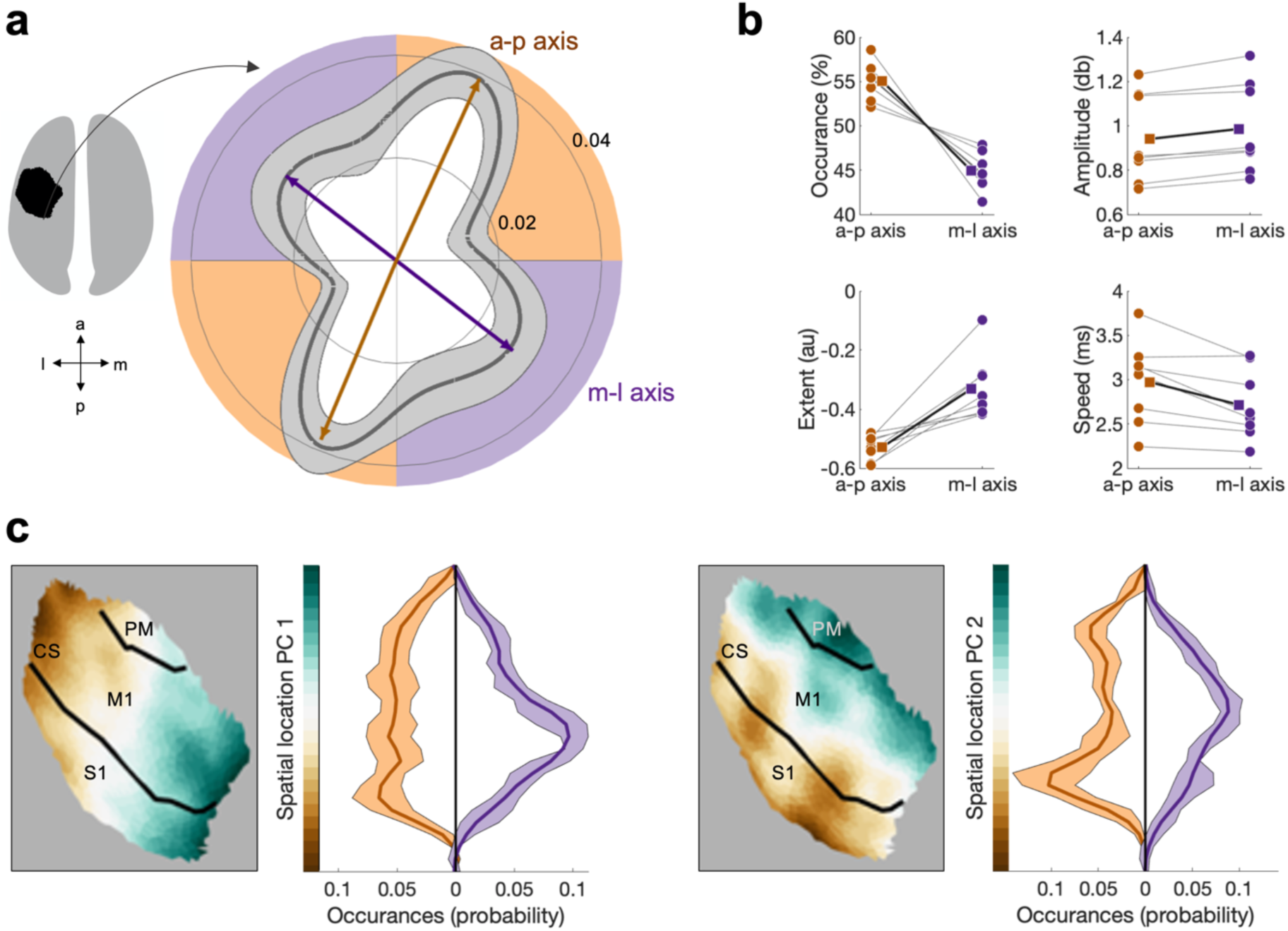
Beta bursts activity propagates along two axes, which have distinct bursts properties. (a) Polar probability histograms showing the probability distribution of burst direction in MNI space. Probability distributions were calculated for each subject individually and then averaged (dark grey line). Variance across subjects is expressed as standard deviation from the mean (light grey area). To estimate the dominant propagation directions, a mixture of von Mises functions was fitted to the averaged probability distribution (arrows). The four functions lie on two axes. One axis has an anterior-posterior orientation which is approximately perpendicular to the orientation of the central sulcus (a-p), while the other axis runs in approximately medial-lateral orientation which is approximately parallel to the orientation of the central sulcus (m-l). (b) Burst occurrence, burst amplitude, burst extent, and burst speed differ as a function of propagation direction. Medians are shown for each subject (circles) and the mean across the subjects’ medians (square). (c) Burst location differs as a function of burst direction. Burst location is described by two Principal Components (PCs) of the Cartesian coordinates of the centre of the burst. For each of the two PCs the surface plot of the component structure and the probability distributions of the PC score are shown. Probability distributions were calculated for each subject individually and then averaged (dark line). Variance across subjects is expressed as standard deviation from the mean (light area). Bursts with a direction parallel to the CS, relative to bursts with a direction perpendicular to the CS, are located more centrally in the ROI. CS, Central Sulcus. S1, Primary Sensory Cortex. M1, Primary Motor Cortex. PM, Premotor Cortex.

The reliability of von Mises functions was assessed using a split-half reliability test. 500 split halves were computed and four von Mises functions estimated on each half independently. The length and direction of the von Mises functions were highly reproducible for all four von Mises functions across both halves of the data (percentage difference in length: *M* = 4.32%, *SD* = 3.86%; angular difference: *M* = 2.2deg, *SD* = 2.8deg; across 500 repetitions and four von Mises functions; **Supplemental Fig. 6**). Further, we tested whether the four von Mises functions were significantly different from zero using non-parametric permutation testing. 5000 permutations were carried out by randomising the propagation direction of each burst and estimating four von Mises functions of the distribution of all bursts. The length of the real van Mises functions were significant while correcting for multiple comparison at *p* < 0.01.

Finally, to examine whether the two main propagation axes can be trivially explained by spatial variability in the beamformer weights, we correlated the latency of the critical points across space before and after regressing out the main components of the spatial variability in the LCMV weights. We found significant correlations (Pearson’s *r: M* = 0.61, *SD* = 0.27 years across individuals, all *p’s* < 0.05), indicating that beamformer weights contribute to, but do not solely explain the observed propagation directions.

Together, these results demonstrates that propagation of sensorimotor beta burst activity propagation occurs along two, orthogonal axes which are oriented approximately parallel and perpendicular the CS.

### Burst characteristics differ as function of the propagation axis

The aforementioned analyses suggest that burst activity propagates along one of two propagation axes. We next asked whether burst propagating along these distinct axes vary in their physiological properties. Specifically, we tested for potential differences in the temporal (i.e., temporal centre), spectral (i.e., frequency centre), or spatial domain (spatial location), as well as burst extent, burst amplitude and propagation speed.

We found significantly more bursts propagating along the a-p axis (*M* = 55.1%, *SD* = 2.0 across individuals), compared to the m-l axis (*M* = 44.9, *SD* = 2.0 across individuals; *T* [test statistic for Wilcoxon signed-rank test, see **Statistical analysis**] = 2.521, *p* < 0.012; **Fig. 4b**). Moreover, bursts propagating along these axes differ in their amplitude, extent, speed (**Fig. 4b**) and spatial location (**Fig. 4c**). Specifically, burst propagating anterior-posterior are characterised by a higher burst amplitude (a-p *M* = 0.94, *SD* = 0.20 across individuals; m-l: *M* = 0.98, *SD* = 0.20 across individuals; *T* = 2.521, *p* < 0.012), larger extent (a-p: *M* = -0.53, *SD* = 0.04 across individuals; m-l: *M* = -0.33, *SD* = 0.1 across individuals; *T* = 2.521, *p* < 0.012) and slower propagation speeds (a-p: *M* = 2.97m/s, *SD* = 0.47m/s across individuals; m-l: *M* = 2.72m/s, *SD* = 0.40m/s across individuals; *T* = 2.38, *p* < 0.017).

The notion that burst activity propagates along distinct anatomical axes was further supported by differences in the spatial location of bursts propagating along these axes. Specifically, the distribution of spatial location PC1 and PC2 (see methods section: **Burst characteristics**; **Supplemental Fig. 7**) differed significantly for burst propagating along axis m-l, relative to bursts propagating along axis a-p for both PC1 (KS [test statistic for Kolmogorov-Smirnov test, see **Statistical analysis**]: *M* = 0.182 across individuals, range = 0.107 – 0.232; 8/8 *p’s* < 0.001) and PC2 (KS: *M* = 0.203 across individuals, range = 0.110 – 0.253; 8/8 *p’s* < 0.001; **Fig. 4c**). This indicates that bursts propagating along a-p are located predominantly in the putative hand region of M1 in the vicinity of the central sulcus, whereas the central locus of burst activity propagating medio-laterally is in S1.

### Distinct physiological fingerprints of pre- and post-movement bursts

Having established that sensorimotor burst activity propagates along two major axes, with distinct foci of burst activation for burst activity propagating along these, we turned to the question whether bursts occurring pre- or post-movement might also be distinguished by their burst and/or propagation properties. To this end, we defined pre-movement bursts as bursts with an on- and offset prior to the movement, and post-movement bursts as bursts with an on- and offset post movement (**Fig. 5a**). Bursts with an onset pre-movement and offset post-movement (*M* = 4.7%, *SD* = 1.9% across individuals) are excluded from this specific analysis. As expected, we found significantly more bursts post- than pre-movement (pre: *M* = 34.3%, *SD* = 5.3% across individuals; post: *M* = 60.9, *SD* = 6.9% across individuals; *T* = 2.521, *p* = 0.012; **Fig. 5b**). Further post-movement bursts are characterised by a larger amplitude (pre: *M* = 0.898db, *SD* = 0.181db across individuals; post: *M* = 0.974db, *SD* = 0.208db across individuals; *T* = 2.521, *p* = 0.012; **Fig. 5b**) and were generally larger in all signal dimensions (burst extent; pre: *M* = -0.719, *SD* = 0.063 across individuals; post: *M* = -0.480, *SD* = 0.036 across individuals; *T* = 2.521, *p* = 0.012; **Fig. 5b**). However, the average spatial location and frequency centre were not significantly different between pre- and post-movement bursts (all *p’s* > 0.1; **Supplemental Fig. 8**).

**Fig. 5:**
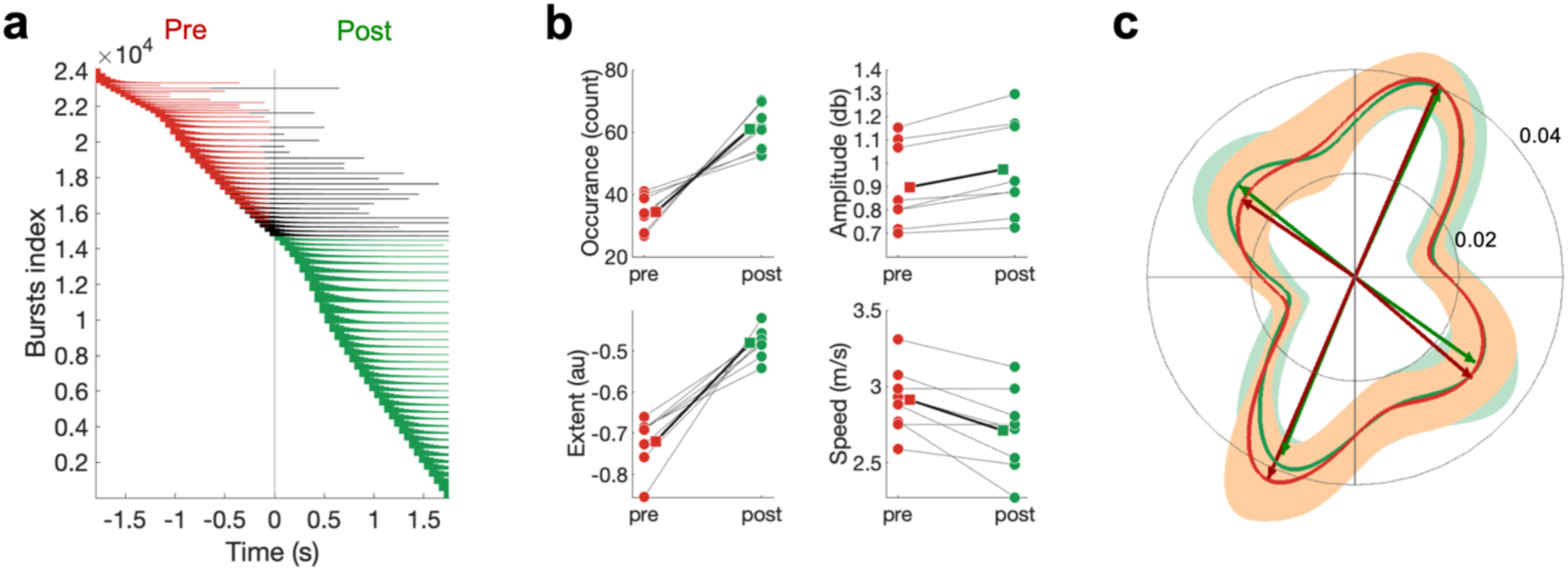
Differences in pre- and post-movement beta bursts. (a) Burst timing relative to the button press across all subjects. Each horizontal line represents one burst. Bursts are sorted by burst onset and burst duration using multiple-level sorting, yielding the burst index. Pre-movement bursts (i.e., bursts that start and end prior to the button press) are highlighted in red, post-movement bursts (i.e., bursts that start after the button press) are highlighted in green, and bursts that start prior to the button press and end after the button press are highlighted in black. (b) Number of bursts, burst amplitude, burst extent and burst speed differ between pre- and post-movement bursts. Medians are shown for each subject (circles) and the mean across the subjects’ medians (square). (c) Propagation direction does not differ between pre- and post-movement bursts. Shown are polar probability histograms separately for pre- and post-movement bursts. Probability distributions were calculated for each subject individually and then averaged (dark line). Variance across subjects is expressed as standard deviation from the mean (light area). Von Mises functions were fitted separately for pre- and post-movement bursts.

Further, in line with non-human primate recordings (Rubino et al., 2006), propagation directions were not significantly different between pre- and post-movement bursts (U^2^ [test statistic for Watson’s U^2^ test, see **Statistical analysis**]*: M* = 0.088 across individuals, range = 0.025 – 0.190; 8/8 *p’s* > 0.1; **Fig. 5c**). The directions of pre-movement bursts activity propagating along the a-p (68/246deg) and m-l direction (148/315deg) did not differ from the directions observed post-movement (a-p: 66/248deg; m-l: 142/325deg). However, while the mean propagation direction did not differ between pre- and post-movement bursts, we found that propagation speed for post-movement bursts was significantly slower than pre-movement (pre: *M* = 2.90m/s, *SD* = 0.20m/s across individuals; post: *M* = 2.69, *SD* = 0.28 across individuals; *T* = 2.521, *p* = 0.012; **Fig. 5b**). Finally, we sought to explore whether pre-movement burst characteristics are related to reaction time. We did not find evidence that burst characteristics relate to reaction time in these data (all *p’s* > 0.1).

## Discussion

The temporal, spectral, and spatial characteristics of beta bursts in human sensorimotor cortex remain unknown. We here show that beta bursts in human sensorimotor cortex occur predominantly post-movement, in the lower beta frequency band, and on the posterior bank of the precentral gyrus. Crucially, sensorimotor beta bursts do not just occur as local standing waves of synchronous activity but propagate along one of two axes that run parallel or perpendicular to the central sulcus, respectively. In addition to the principal axis of their propagation direction, these bursts differ in their occurrence, location, propagation speed, amplitude, and extent. Further, post-movement bursts are characterised by higher amplitude, larger extent and are slower propagation speed, suggesting distinct physiological markers and functional roles pre- and post-movement. However, the comparable spectral and spatial centres as well as the propagation direction of pre- and post-movement bursts indicate the same underlying burst generator. Collectively, our data provide novel evidence that a substantial proportion of human sensorimotor beta burst activity travels along two anatomical and functional distinct axes, with distinct burst properties pre and post movement.

### Distinct anatomical propagation axes of sensorimotor beta activity

Traveling wave activity occurs at multiple spatial scales, ranging from mesoscopic, columnar to macroscopic, transcortical levels (Muller et al., 2018). Here we show that beta burst activity in human sensorimotor cortex propagates along two approximately orthogonal axes that are oriented in anterior-posterior and medial-lateral direction. Recordings from invasive multi-electrode arrays confirm the dominance of these two propagation axes, albeit on a smaller spatial scale of roughly 4mm. For example, Takahashi and colleagues reported that beta activity in M1 of a tetraplegic patient propagated along the medial-lateral axis (Takahashi et al., 2011). In non-human primates, beta activity propagates along the anterior-posterior axis in M1 (Balasubramanian et al., 2020; Best et al., 2016; Rubino et al., 2006; Takahashi et al., 2011, 2015), and along the medial-lateral axis in the dorsal premotor cortex (Rubino et al., 2006), indicating regional differences in spatiotemporal patterns (Rubino et al., 2006; Rule et al., 2018).

In these studies, neural activity has been recorded from a single cortical region, limited by the dimension of the electrode array (roughly 0.16 cm^2^). By contrast, we here identified spatiotemporal patters of beta activity in burst events of an average apparent spatial width of ∼6cm^2^ located in M1 and adjacent cortical areas. By leveraging high SNR MEG recordings that permit high sensitivity in all signal domains, we were able to quantify bursts and their spatiotemporal pattern non-invasively over these functionally cogent brain regions at a spatial scale that sits between invasive recordings in animals and previous human M/EEG or intracranial recordings (Alexander et al., 2016; Roberts et al., 2019; Rule et al., 2018; Stolk et al., 2019; Takahashi et al., 2011). Our results extend previous invasive recordings by showing that bursts activity can travel across sensory and motor cortices and bridge across functionally distinct brain areas.

The spatial profiles of propagation of beta activity along the anterior-posterior and medial-lateral direction are in line with the idea that propagation directions are imposed by the dominant internal connections within anatomical networks (Rubino et al., 2006). Here, our dominant propagation axes conformed to an anterior-posterior network comprising dorsal premotor cortex, primary motor cortex and primary sensory cortex (Cauller et al., 1998; Kurata, 1991; Luppino & Rizzolatti, 2000; Muakkassa & Strick, 1979) and a medial-lateral network spanning across medial and lateral dorsal premotor cortex, supplementary motor area cortex and caudal portions of ventral premotor cortex (Dum, 2005; Ghosh & Gattera, 1995; Luppino et al., 1993). The latter is thought to mirror proximal and distal sites within the motor cortex (Rubino et al., 2006), with proximal representations (i.e., shoulder and elbow) located more medially and distal representations (i.e., wrist and fingers) located more laterally (Penfield & Boldrey, 1937). This suggests that at a macro-scale level, the direction of wave propagation is dictated by the underlying horizontal connections, though further work across different spatial scales (such as (Sreekumar et al., 2020)) is required to fully unpack the precise relationship between sustained rhythmic synchronous spiking activity within neural populations, mesoscopic and macroscopic traveling wave activity.

While our results further corroborate the importance of anterior-posterior and medial-lateral propagation axes, the precise mechanism of travelling wave activity remains unclear. One possible mechanism is that excitation from a single generator propagates through a network, guided by conduction delays within corticocortical and the corticothalamic system (Ermentrout GB, 2001; Muller et al., 2018; Prechtl et al., 2000). Alternatively, travelling wave activity could arise from one generator driving a network through increasing time delay, so-called fictive traveling waves, or coupled generators that exhibit stable phase differences. Different levels of network interactions may thus generate and sustain propagating waves. Common to all travelling wave activity is the idea that they generate a consistent spatiotemporal frame for further neuronal interactions. In mesoscopic data it is very challenging to analytically resolve any ambiguity about the mechanism of wave generation. LFP recordings with implanted electrode arrays in non-human primates suggest that coupled oscillators contribute significantly to beta travelling waves over a spatial scale of 0.16cm^2^ (Rule et al., 2018).

### Propagation axes of sensorimotor beta activity are physiologically distinct

While previous work has investigated individual aspects of neural activity in relation to propagation direction (Balasubramanian et al., 2020; Bhattacharya et al., 2022) we here consider all signal domains of neural activity. We found that the two propagation axes can be distinguished based on their physiological properties, such as propagation speed, burst occurrence, amplitude, and extent. Specifically, beta activity propagating in the medial-lateral direction is characterised by higher burst amplitude and larger burst extent, i.e., bursts are larger in all signal domains. Further, more bursts propagate along the anterior-posterior l direction, which is also characterised by faster propagation speed.

Propagating wave activity can occur in a wide range of different speeds, with propagation speeds broadly falling into two categories. Speeds for mesoscopic traveling waves occurring within cortical columns and their lateral connections, as identified using local field potential (LFP), multielectrode arrays or optical imaging, and range between 0.1-0.8m/s (Bhattacharya et al., 2022; Rubino et al., 2006; Takahashi et al., 2011, 2015). These slower wave speeds are consistent with axonal conduction speeds of unmyelinated horizontal fibres in the superficial layers of the cortex (Girard et al., 2001).

By contrast, macroscopic traveling waves spanning across several cortical regions, and commonly assessed using mass-neural signal recordings such as M/EEG or ECoG, have been reported at speeds ranging from around 1-10m/s (Alexander et al., 2016; Hughes, 1995; Muller et al., 2018). The relatively large variability in propagation speed of macroscopic traveling waves is partly due to variability in spatial resolution with low spatial resolution being susceptible to aliasing artefacts (Alexander et al., 2016; Bahramisharif et al., 2013), and some uncertainty in the travelled distance. Regarding the latter, while it has been recommended to study travelling waves on the cortical surface (Alexander et al., 2019; Hughes, 1995) it is still unclear whether neural activity truly propagates along the brains cortical surface (as quantified by geodesic distance), or, at least in part, propagate through the brain volume (as quantified by Euclidian distance). Further, propagation distance can be computed on the original, folded cortical surface or on the inflated surface. Our data show that propagation speed derived from the folded surfaces is roughly twice as fast than the propagation speed derived from the inflated surface (**Supplemental Fig. 9**), which is in line with the previously reported folding factor of x2.2 (Alexander et al., 2016; Burkitt et al., 2000). Notwithstanding the uncertainty this introduces in estimating propagation speeds, the range of propagation speeds observed here are compatible with previous reports from human and non-human primates (Hughes, 1995; Muller et al., 2018), and are compatible with axonal conduction speeds of myelinated cortical white matter fibres (Swadlow & Waxman, 2012), suggesting an active role for macro-scale traveling burst activity in intra-areal communication and information transfer.

### Pre- and post-movement burst share the same generator expressed differently

The transient bursts of beta activity in our human MEG data lasted, on average, several hundred milliseconds, and span over approximately 3 Hz predominantly in the lower beta frequency range. These temporal and spectral properties are broadly in line with previous reports (Cagnan et al., 2019; Quinn et al., 2019; Seedat et al., 2020; Shin et al., 2017; Sporn et al., 2020; Tinkhauser, Pogosyan, Little, et al., 2017), with variation in the absolute values being strongly dependent on how bursts are operationalised (Zich et al., 2020). We extend these previous reports on the temporal and spectral burst characteristics, by additionally characterising spatial burst characteristics. Sensorimotor beta burst activity often spans over several square centimetres with a distinct topographic distribution. The majority of bursts are located on the posterior bank of the precentral gyrus, with a proportion of bursts that spread to adjacent areas. While approaching the spatial limits of human MEG, these data indicate the possibility of locating beta activity within the sensorimotor cortex.

To further elucidate the generator processes and functional roles of sensorimotor beta bursts we next compared pre- and post-movement bursts with regard to both their temporal, spectral and spatial burst characteristics, and their propagation properties. We confirmed that post-movement, compared to pre-movement, bursts occur more frequently, and are stronger (i.e., higher burst amplitude) and larger in all signal domains (i.e., larger burst extent). These observations are largely in line with previous studies (Quinn et al., 2019; Seedat et al., 2020; Zich et al., 2018), whereby we note that (Little et al., 2019) no difference in temporal burst duration between pre- and post-movement bursts was reported. We believe this discrepancy is because (Little et al., 2019) employed different thresholds for pre- and post-movement bursts, whereby here the same threshold was used. Moreover, we find that pre-movement bursts exhibit faster propagation speed than post-movement burst activity. There is no evidence that the difference in propagation speed is mediated through differences in the frequency (Alexander et al., 2016), or spatial location of bursts, as both metrics are comparable for pre- and post-movement bursts. The functional relevance of this difference in propagation speed merits further consideration in the future, but it indicates that parsing the functional role of beta activity may require its decomposition into its physiologically distinct stationary and propagating components. Finally, we show that pre- and post-movement bursts propagate along the same propagation axis, which is in line with previous reports, observing the same propagation axes during action (Rubino et al., 2006) and rest (Takahashi et al., 2011). This provides further evidence that the propagation of burst activity is constrained by the underlying connectivity.

Together, our results show that, compared to pre-movement bursts, post-movement bursts are stronger and larger in all signal domains, whereby their spectral and spatial centre, as well as their propagation direction, are comparable. We believe this indicates that pre- and post-movement bursts, therefore, share the same generator processes, which exhibits more and stronger bursts post-movement. Based on biophysical principled neural modelling, corticocortical and thalamocortical circuit mechanisms are thought to play a critical role in generating sensorimotor beta bursts (Bonaiuto et al., 2021; Neymotin et al., 2020; Law et al., 2022). Interestingly, sensorimotor beta bursts have not only been observed during action but also during rest (Zich et al., 2018; Seedat et al., 2020; Becker et al., 2020; Echeverria-Altuna et al., 2021), which raises the question of their functional role. That sensorimotor beta bursts occur across functional states, spatial scales and species suggests that the functional role of the mere presence of bursts is a very elementary one, such as maintaining the ‘status-quo’ (Engel & Fries, 2010) or ‘null space’ (Kaufman et al., 2014). In addition, we believe that specific functional roles can be linked to the manifestation of bursts quantifiable by their temporal, spectral and spatial bursts characteristics as well as their propagation properties. To give one example, motor symptoms in Parkinson’s disease have been linked to prolonged burst duration (Deffains et al., 2018; Tinkhauser, Pogosyan, Little, et al., 2017; Tinkhauser, Pogosyan, Tan, et al., 2017). The proposed hierarchical dual-role framework of burst function can be tested using biophysical models (Neymotin et al., 2020) and targeted neuromodulatory experiments.

### Caveats of spatial and spatiotemporal properties in source space

Non-invasive techniques have limitations that should be considered when interpreting the spatial domain of bursts and travelling wave activity. LCMV beamformers assume that each source is a single dipole and that there are no other correlated sources in the brain. These limitations make interpretation of spatial structure in LCMV power maps ambiguous. We explore several specific issues: firstly, whether the apparent spatial extent of a source is simply modulated by the SNR of the signal. Secondly, the inherent smoothness of the source reconstruction maps due to the mapping of a few hundred sensors to several thousand voxels. Finally, if a patch of cortex is active rather than a single point-source, then these correlated voxels can suppress the signal of interest. Each of these points can be challenging when interpreting the spatial domain of bursts and travelling waves.

The first issue suggests that differences in the bursts’ apparent spatial width could simply be caused by differences in SNR across and/or within sessions rather than differences in the spatial distribution of cortical activity. We performed one beamformer per session, thus different SNR levels across sessions would affect the beamformer weights. However, if variation across sessions in beamformer weights would explain variation in bursts’ apparent spatial width, we would expect a negative relationship between burst amplitude and burst apparent spatial width across sessions. This is not the case in our data, suggesting that between-session differences in beamformer weights do not cause the observed differences in bursts’ apparent spatial width. Nevertheless, spatial width of burst activity measured with M/EEG or ECoG should be interpreted with caution. Here, due to the strong correlation between the bursts’ apparent spatial width, temporal duration, and frequency spread, we combined these signal properties using PCA and used the resulting cross-modal measure burst extent.

Secondly, the inherent smoothness of the beamformer solution can lead to ‘trivial’ structure in the source solution, meaning that single sources can leak across cortex or that multiple sources can become blurred together. Across space in bursts diverse phase lags exists suggesting that structure is unlikely to have arisen solely from leakage of a single source. The functional role of travelling waves remains unclear. As outlined above, the mechanisms underlying travelling waves remain ambiguous (see discussion section: **Distinct anatomical propagation axes of sensorimotor beta activity**), both at the meso- and macro scale (Hughes, 1995; Muller et al., 2018). We cannot rule out the possibility that this phase structure arises from mixing of multiple distinct sources but take a ‘gradient’ or ‘travelling wave’ perspective here to better link with comparative literature. While this concerns travelling wave analyses across a range of spatial scales and recording techniques; source space analysis, as employed here, entails an additional issue – namely whether the propagation directions can be trivially explained by spatial variability in the beamformer weights. Our control analysis showed that the estimated propagation direction correlates significantly with the propagation direction obtained after regressing out the main components of spatial variability in the beamformer weights. This indicates that beamformer weights can contribute to, but do not solely explain spatiotemporal gradients in human MEG data.

Finally, patches of high amplitude, correlated sources can be mutually suppressed by the LCMV beamformer leading to an apparent loss of signal. Though we cannot remove the possibility these mutual correlations may be suppressing part of the signal, we observe strong task-related activity suggesting that a substantial proportion remains in our analysis.

Together, we acknowledge that the beamformer weights can affect bursts’ spatial width and propagation direction but believe that our control analyses suggests that the beamformer weights are not driving the observed effects.

## Materials ad Methods

### Participants and experimental task

The study was approved by the UCL Research Ethics Committee (reference number 5833/001) and conducted in accordance with the Declaration of Helsinki. Informed written consent was obtained from all participants. All participants (6 male, *M* = 28.5 years, *SD* = 8.52 years across individuals) were free of neurological or psychiatric disorders, right-handed and had normal or corrected-to-normal vision.

Participants performed a visually cued action decision making task in which they responded to visual stimuli projected onto a screen by pressing one of two buttons using their right index or middle finger (for details see (Bonaiuto et al., 2018)). The task uses a factorial design with congruence (congruent, incongruent) and coherence (low, medium, high). Here we only consider congruent, high coherence trials (42 trials per block) that were responded to correctly (for full design see (Bonaiuto et al., 2018; Little et al., 2019)).

### MRI acquisition and processing

Prior to the MEG sessions, structural MRI data were acquired using a 3T Magnetom TIM Trio MRI scanner (Siemens Healthcare, Erlangen, Germany). A T1-weighted 3D spoiled fast low angle shot (FLASH) sequence was acquired to generate an accurate image of the scalp for head-cast construction. Subject-specific head-casts optimise co-registration and reduce head movements, and thereby significantly improve the signal to noise ratio. See (Bonaiuto et al., 2018; Meyer et al., 2017; Troebinger et al., 2014) for details on the sequence and the head-cast construction.

In addition, a high-resolution, quantitative, multiple parameter mapping (MPM) protocol, consisting of 3 differentially-weighted, RF and gradient spoiled, multi-echo 3D FLASH acquisitions recorded with whole-brain coverage at 800 mm isotropic resolution, was performed. See (Bonaiuto et al., 2018) for details on the protocol. Each quantitative map was co-registered to the scan used to design the head-cast, using the T1 weighted map. Individual cortical surface meshes were extracted using FreeSurfer (v5.3.0; (Fischl, 2012)) from multiparameter maps using the PD and T1 sequences as inputs, with custom modifications to avoid tissue boundary segmentation failures (Carey et al., 2018). Meshes were down-sampled by a factor of 10 (vertices: *M* = 30,095, *SD* = 2,665 across individuals; faces: *M* = 60,182, *SD* = 5,331 across individuals) and smoothed (5mm). Here we used the original and the inflated pial surface.

### MEG acquisition and pre-processing

MEG data were acquired using a 275-channel Canadian Thin Films (CTF) MEG system using individual head-casts in a magnetically shielded room. Head position was localised using three fiducial coils placed at the nasion and left/right pre-auricular points, within the head-cast. Data were sampled at 1200Hz. A projector displayed the visual stimuli on a screen (∼50 cm from the participant), and participants made responses with a button box.

A summary of the data processing pipeline is shown in **Supplemental Fig. 1**. MEG data were analysed using the OHBA Software Library (OSL: https://ohba-analysis.github.io/osl-docs/). MEG data were processed in for each block separately unless stated otherwise. Firstly, raw data were converted to SPM12 format for analysis in Matlab2019b. Registration between structural MRI and the MEG data was performed with RHINO (Registration of head shapes Including Nose in OSL) using only the Fiducial landmarks and single shell as forward model. Unless stated otherwise data were analysed in single subject space.

Continuous data were down-sampled to 300Hz. Further, a band-pass (1-95Hz) and notch-filter (49-51Hz) were applied. Time segments containing artefacts were identified by using generalised extreme studentized deviate method (GESD (Rosner, 1983)) on the standard deviation of the signal across all sensors in 1s non-overlapping windows, with a maximum number of outliers limited to 20% of the data and adopting a significance level of 0.05. Data segments identified as outliers were excluded from subsequent analyses.

Further, denoising was applied using independent component analysis (ICA) using temporal FastICA across sensors (Hyvarinen, 1999). 62 independent components were estimated and components representing stereotypical artefacts such as eye blinks, eye movements, and electrical heartbeat activity were manually identified and regressed out of the data.

Data were then filtered to the frequency band of interest (β 13-30 Hz) and segmented from - 2s to 2s relative to the button press. Segmented data were projected onto subjects’ individual cortical surface meshes using a Linearly Constrained Minimum Variance (LCMV) vector beamformer (Van Veen & Buckley, 1988; Woolrich et al., 2011). The beamformer weights were estimated at the centre of each face, referred to henceforth as spatial locations. A covariance matrix was computed across all segments and was regularised to 55 dimensions using principal component analysis (PCA). All analyses are conducted in source space.

### Time-frequency decomposition

Time-frequency analysis was applied to single trials and spatial locations using dpss-based multitaper (window = 1.6s, steps = 200ms) with a frequency resolution of 1Hz. Epochs were baseline corrected (−1.8s to -1.1s). This procedure results in a trial-by-trial time-frequency decomposition for each spatial location, i.e., relative power in 4D, time x frequency x space x trial, whereby space is on its own 3-dimensional (x, y, z coordinates of surface locations).

### Burst operationalisation

We used binarization and high-dimensional clustering to operationalize beta bursts. Power derived from time-frequency analysis was first binarized using a simple amplitude threshold (see **Supplemental Methods**). The threshold was obtained empirically, as in previous work (Little et al., 2019). Specifically, trial-wise power was correlated with the burst probability across a range of different threshold values (median to median plus seven standard deviations, in steps of 0.25). The threshold value that retained the highest correlation between trial-wise power and burst probability was used to binarize the data. To account for difference in signal-to-noise across sessions, days and subjects we obtained one threshold per session (*M* = 2.97 x SDs above mean, *SD* = 0.66 across sessions; **Supplemental Fig. 2**).

Following binarization, data were clustered across time, frequency, and space on a single trial level (see **Supplemental Methods**). Burst identification was limited to the time of interest (−2 to 2s relative to the button press), the frequency of interest (13-30Hz) and region of interest (ROI, left-hand area). To restrict the burst analysis to a ROI, volume-based ROIs in MNI space were normalised to subject’s native space using the inverse deformation field and transformed to surface-based ROIs. Clusters had to span at least 2 time points, frequency steps and spatial locations to be considered further.

### Burst characteristics

We divide burst characteristics into 1^st^ and 2^nd^ level burst characteristics. We define 1^st^ level burst characteristics as characteristics that are obtained for each burst and each domain separately. **Fig. 1** illustrates the first level characteristics. For the temporal domain, burst temporal on- and offset, temporal duration and centre (i.e., mean of on- and offset) were obtained. Equally, low and high frequency boundaries, frequency spread and centre (i.e., mean of low and high boundary) were extracted for the spectral domain. For the spatial domain we obtained the spatial width (i.e., total surface area defined as the sum of the area of all faces), the size in each dimension (x, y, z) using the minimum bounding rectangle (i.e., bounding box), and the spatial centre. The spatial centre is defined as the projection of the centre of mass onto the surface. The spatial centre can be described using its Cartesian coordinates. An alternative to the description of the spatial centre is provided by the first two components of a PCA of the Cartesian coordinates (**Supplemental Fig. 7**). The first two PCs describing 98% of variance are retained for further analysis. The first PC (76.3% variance explained) contains a spatial gradient along the anterior/lateral – posterior/medial axis. The second PC (22.3% variance explained) contains a spatial gradient along the anterior/medial – posterior/lateral axis. Thus, the location of an individual burst can be described by the two PC scores, relating to the amount of each PC that it contains. In addition, burst amplitude was obtained, i.e., the mean amplitude across all time points, frequencies, and spatial locations of the burst.

These 1^st^ level burst characteristics form the basis for 2^nd^ level burst characteristics. These can be broadly summarised as a) combinations and b) interactions of characteristics within and across domains. Here we extract one of these measures: temporal duration, frequency spread, and apparent spatial width were combined to a single metric, i.e., burst extend. The three measures are highly correlated within subjects (*M* = 0.785; *SD* = 0.021, across the three correlations and eight individuals, **Supplemental Fig. 10a,b**) and where therefore reduced to a single metric using PCA. The first principal component explains 85.6% of the variance and is defined as burst extent (PC 2: 7.5%, PC 3: 6.9%, **Supplemental Fig. 10c**).

### Propagation direction and speed of neural activity within bursts

To investigate whether activity within human sensorimotor bursts propagates, we identified the dominant propagation direction and speed for each burst. To this end, data (before time-frequency decomposition, see **Supplemental Fig. 1**) of each burst were extracted from burst on- to offset for each surface location in the burst. The sign ambiguity in the beamforming process entails that the spatial locations within a burst may have arbitrarily opposite signs. This is not an issue when estimating power, as above, but can impact on the estimation of the propagation direction. Sign ambiguity was resolved using the sign-flipping algorithm described in (Vidaurre et al., 2018). For a finer temporal resolution data were interpolated by a factor of 10.

For each burst, we estimated the propagation direction and propagation speed. Propagation direction and speed were estimated from critical points in the oscillatory cycle (four critical points per oscillatory cycle, i.e., peak and trough as well as peak-trough and trough-peak midpoint, grey vertical lines in **Fig. 2a**) and then averaged across critical points within one burst.

The propagation direction at each critical point was estimated from the relative latency (i.e., absolute latency of that critical point at each surface location relative to absolute latency of that critical point for the average across all surface locations in that burst). For example, **Fig. 2b_i_** shows the relative latency for each surface location in the burst for the critical point at 212ms in the burst. Next, from these relative latencies and their surface locations the propagation direction was estimated. Specifically, propagation direction was estimated using linear regression (Balasubramanian et al., 2020), whereby the relative latency at the surface location was predicted from the coordinates of the surface location of the inflated surface. On the inflated surface, a gradient in the z-direction is always depicted by a gradient in x- or y-direction, which is why only two simple linear regressions were estimated, one for the x- and one for the y-direction (**Fig. 2b_ii_**). Propagation direction along the x-y-direction was obtained by transforming the regression coefficients from Cartesian coordinates to spherical coordinates (red arrow in **Fig. 2b_iii_**). For each regression, its associated coefficient of determination (R^2^) was calculated and the two R^2^’s averaged. This approach results in one propagation direction and one R^2^ per critical point.

Propagation direction across critical points was obtained by clustering (i.e., Spectral clustering) the propagation directions across critical points. Three scenarios existed: 1) One cluster was obtained and the variance across directions of critical points was relatively low (standard deviation < π/4; **Fig. 2b_iii_**; **Supplemental Fig. 5a**); 2) One cluster was obtained and the variance across directions of critical points was relatively high (standard deviation > π/4; **Supplemental Fig. 5b**); 3) More than one cluster was obtained (**Supplemental Fig. 5c**). Scenario 2 and scenario 3 indicate complex propagation patterns, such as random or circular patterns (Denker et al., 2018; Rule et al., 2018). Based on previous literature we expect planar traveling waves to be dominant in the primary motor cortex (Balasubramanian et al., 2020; Rubino et al., 2006; Rule et al., 2018; Takahashi et al., 2011). For bursts of scenario 1 propagation directions and R^2^ were averaged across critical points (back arrow in **Fig. 2b_iii_**). To have sufficient confidence in the direction bursts with an average R^2^ < 0.2 were discarded (Balasubramanian et al., 2020). Following this procedure, we found that 79.59% (*SD* = 2.37% across individuals) of the bursts show a spatiotemporal pattern (R^2^: *M* = 0.35, *SD* = 0.02 across individuals). To combine propagation directions across subjects, propagation directions were spatially normalised to MNI space using the deformation field. Directions are presented as probability distributions. On the average of the probability distributions across subjects the propagation direction was quantified using a mixture of von Mises functions.

The propagation speed at each critical point was defined as the distance between the spatial locations with the largest and smallest relative latency divided by the difference in their latencies (Bahramisharif et al., 2013). Distance was computed using exact geodesic distance (**Fig. 2c_ii_**) on the inflated surface. See **Supplemental Methods** and **Supplemental Fig. 9** for a comparison of propagation speed when computing the distance on the inflated surface and the original surface. Propagation speed was averaged across critical points.

### Accuracy of the propagation direction detection in simulated and real meshes

Using simulation, we evaluated the accuracy of the propagation direction estimation. To this end, we generated 360 noise-free high-resolution gradients span 1deg in steps of 1deg (**Fig. 3a** shows a subset). To evaluate the effect of mesh type and spatial sampling we created three 2D mesh types, 1) square mesh (**Fig. 3b**), 2) circular mesh (**Fig. 3c**), and 3) random mesh (**Fig. 3d**), whereby each mesh type was sampled at three spatial sampling rates: N/2, N, and Nx2 (N approximates the spatial sampling of the surface mesh, i.e., roughly 27 surface locations per cm^2^). For each gradient and each mesh, the propagation direction was estimated and the estimation error, i.e., difference between true and estimated propagation direction, computed. For the random mesh, this procedure was repeated 100 times, each time with a different random mesh.

As the surface mesh is irregular and each burst is unique in its spatial size and shape, we additionally evaluated the accuracy of the propagation direction estimation for the real bursts. To this end, for each individual burst and each gradient, the propagation direction was estimated, and the estimation error computed as above.

### Control analysis

The ill-posed nature of the inverse problem in M/EEG means that the source estimation has a degree of smoothness. While this is unavoidable and shared with all inverse problem methods, the smoothness can be problematic when interpreting the spatial domain of burst and their spatiotemporal gradients, travelling waves. We perform a series of control analysis to explore the practical effect of these ambiguous in our data. Our reasoning was that with regards to interpreting the spatial width of burst activity, any differences could be caused by differences in SNR across and/or within sessions rather than differences in the spatial distribution of cortical activity (see **Supplemental Fig. 4ai, bi** for a schematic illustrations). To address this, we performed several correlation analyses between burst amplitude and burst apparent spatial width, between and across sessions.

Regarding the interpretation of traveling waves, there is inherent ambiguity concerning the mechanisms that generate a travelling wave (see discussion section: **Distinct anatomical propagation axes of sensorimotor beta activity**; Prechtl et al., 2000; Ermentrout and Kleinfeld 2001). While this concerns travelling wave analyses across a range of spatial scales and recording techniques, the source space analysis employed here entails an additional issue – namely whether the propagation directions can be trivially explained by spatial variability in the LCMV weights. To address this issue, we correlated the latency of the critical points across space before and after regressing out the main components of the spatial variability in the LCMV weights. Specifically, we performed PCA on the LCMV weights and retained the components that explained 90% of the variance in the LCMV weights. We then performed, for each critical point of each burst, a multiple regression analysis with the latencies of the critical point across space as dependent variable and the coefficients of the PCs across space as independent variables. We then correlated the latency of the critical points across space with the residuals of the multiple regression. Pearson’s *r* was first averaged across critical points within bursts, and then across bursts.

### Statistical analysis

Statistical analysis was performed using nonparametric testing in Matlab2019b. If not stated otherwise, descriptive statistics depict mean and standard deviation of the median across subjects. Burst characteristics with unimodal distributions (e.g., burst amplitude, burst propagation speed), were compared using Wilcoxon signed-rank test on the medians of the distribution. The test statistic is reported as a value of *T*. Burst characteristics with multimodal distributions (e.g., spatial location) were compared using two-sample Kolmogorov–Smirnov test on the single subject level. Test statistic is reported as value of *KS* (i.e., mean and range across subjects). Two circular distributions (e.g., propagation direction pre- and post-movement) were compared using two-sample Watson’s U^2^ test (Landler et al., 2021) on the single subject level. Test statistic is reported as value of *U^2^* test (i.e., mean and range across subjects).

To test whether there is significant spatiotemporal structure in burst activity, we compared the propagation direction of real and surrogate data. Specifically, for a subset of bursts, i.e., 100 randomly selected bursts, 1000 surrogates were created for each burst from the data after sign ambiguity was resolved (see **Supplemental Fig. 1**). Surrogate data were obtained by computing the discrete Fourier transform of the data, randomizing the phase spectrum while preserving the amplitude spectrum, and then computing the inverse discrete Fourier transform to obtain the surrogated data (method 3 in (Hurtado et al., 2004)). For each burst, the magnitude of the propagation direction of the real data was compared to the distribution from 1000 surrogates.

To quantify the overall propagation direction, a mixture of four von Mises functions was fitted to the average of the subjects’ probability distribution of propagation directions across bursts. This provides an estimate of the angle and length of the von Mises functions. Reliability of von Mises functions was assessed using a split-half reliability. 500 split halves were computed and four von Mises functions estimated on each half independently. For both, angle and length, the difference between the two halves was computed. Further, to test whether the von Mises functions were significantly different from zero, non-parametric permutation testing was employed on the length of the von Mises functions. Permutations were carried out by randomising the propagation direction of each burst. 5000 permutations were computed before statistical significance was determined on the length of the von Mises functions while correcting for multiple comparison at p<0.01.

## Acknowledgements

C.Z. is supported by the Brain Research UK (201718-13, 201617-03). A.J.Q. is supported by the NIHR Oxford Health Biomedical Research Centre and a Wellcome Trust Strategic Award (098369/Z/12/Z). G.C.O. is supported by EPSRC (EP/T001046/1) funding from the Quantum Technology hub in sensing and timing (sub-award QTPRF02). L.C.M. is supported by the Medical Research Council (MR/N013867/1). The Wellcome Centre for Human Neuroimaging and The Wellcome Centre for Integrative Neuroimaging are supported by core funding from the Wellcome Trust (203147/Z/16/Z and 203139/Z/16/Z, respectively).

## Supplemental Information

### Supplemental Methods

#### Burst threshold

Detecting bursts simultaneously in the temporal, spectral and spatial domain is accompanied by some conceptual and computational challenges. Here we opt for a simple thresholding approach, rather than a more data driven approach, such as the Hidden Markov Model (HMM, (Quinn et al., 2019; Vidaurre et al., 2016)). Firstly, existing HMM variants do not provide the here desired frequency resolution. Secondly, adapting the amplitude-envelope HMM to threshold power derived from time-frequency analysis poses a computational challenge for this high-dimensional dataset. Finally, one of the main advantages of HMM, i.e., the prevention of burst-splits (see (Quinn et al., 2019)), is overcome in the 5D clustering procedure. Together, while HMM, and other data driven approaches are generally advantageous, in this framework a simple amplitude threshold is preferred.

Another aspect worth highlighting affects the threshold to detect bursts, which can be summarised as follows: Are bursts better binarised by using a uniform or adaptive threshold across time, frequency, and space? Here we opt for the former approach, as it allows for direct comparisons across different points in time, frequency, or space. On the other hand, the latter has the potential of accounting for differences in SNR across time, frequency, and space.

#### 5D clustering

To obtain 3D bursts, binarized data were clustered using a three-stage approach (see **Supplemental Video 1**). Note that data are 4-dimensional, i.e., time x frequency x space x trial, whereby space is on its own 3-dimensional (x, y, z coordinates of surface locations). First, for each trial data were clustered in 2D (i.e., time x frequency). To this end, the binarized data were summed over the spatial domain and time-frequency cells with at least one surface location being ‘on’ were clustered using 8-connectivity (i.e., connected horizontally, vertically, or diagonally).

Second, for each time-frequency cell with at least one surface location being ‘on’, spatial locations on the surface mesh were clustered in 3D (i.e., x, y, z coordinates of surface locations). Spatial locations were part of the same cluster if their Euclidean distance was smaller than the maximal distance of two spatial locations (*M* = 2.66mm; *SD* = 0.15mm across individuals).

Finally, source clusters were combined across time-frequency cells using 8-connectivity, i.e., if two spatial clusters of two adjourning time-frequency cells overlapped in at least one surface location the two spatial clusters were combined. This procedure allows clustering in high-dimensional irregular space and results in 3D (time x frequency x space) bursts.

#### Propagation speed

Propagation speed was calculated by dividing the distance between the spatial locations with the largest and smallest relative latency (i.e., latency of each surface location relative to the average latency) by the difference in their latencies (Bahramisharif et al., 2013). Distance can be computed either on the original surface or on the inflated surface (**Supplemental Fig. 9**). The speed computed using the distance on the original surface (*M* = 4.90m/s, *SD* = 0.46m/s across individuals) is faster than the speed computed using the distance on the inflated surface (*M* = 2.61m/s, *SD* = 0.39m/s across individuals). This difference is well in line with a suggested cortical folding factor of x2.2 to adjust propagation speeds for cortical folding (Alexander et al., 2016; Burkitt et al., 2000). Propagation speed is in the expected range of macroscopic waves (Hughes, 1995; Muller et al., 2018).

---

***Supplemental Video 1***

*Same as Fig. 2, but each frame corresponds to a different critical point within the burst*.

**Supplemental Fig. 1.**
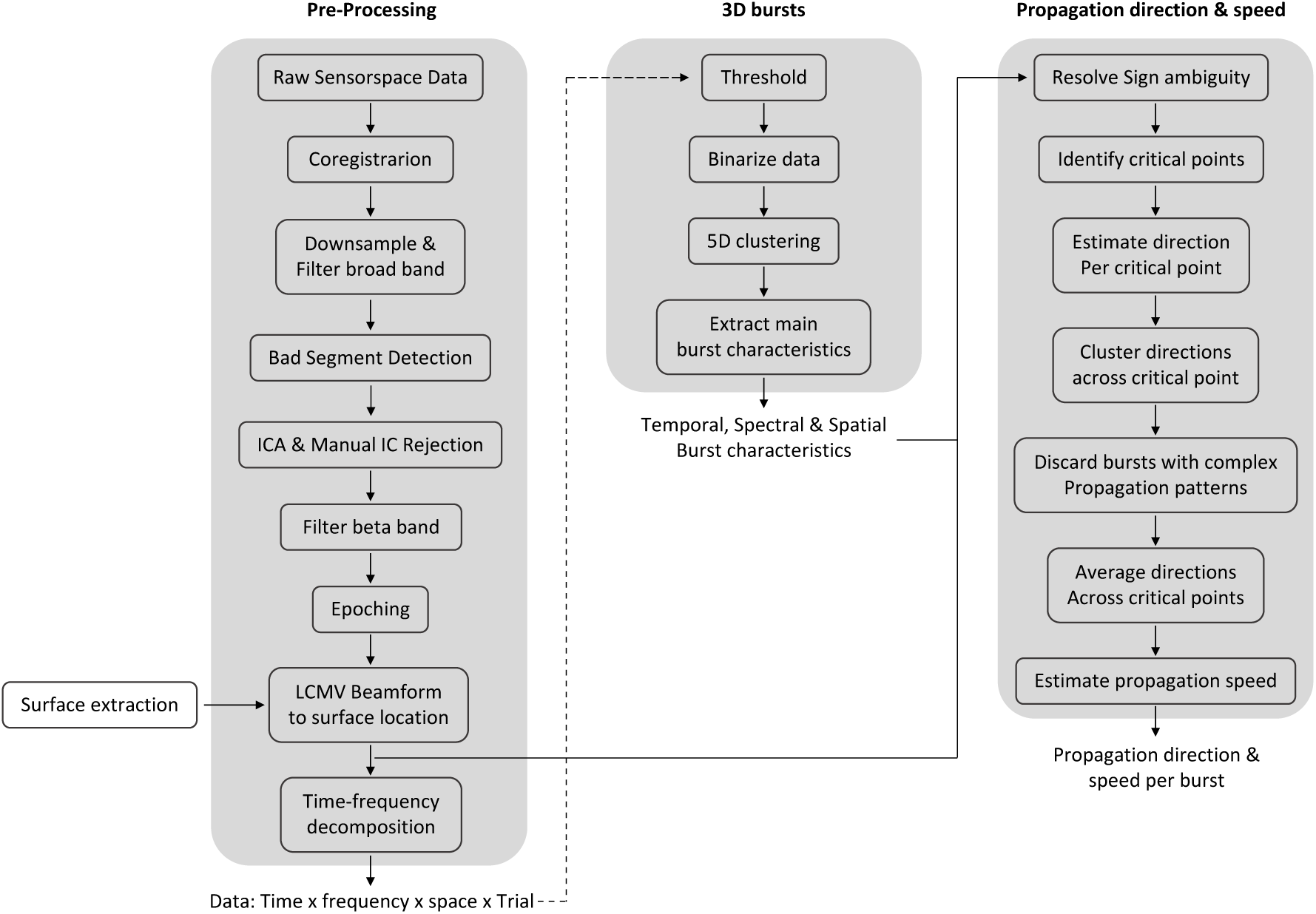
A schematic for the processing pipeline.

**Supplemental Fig. 2.**
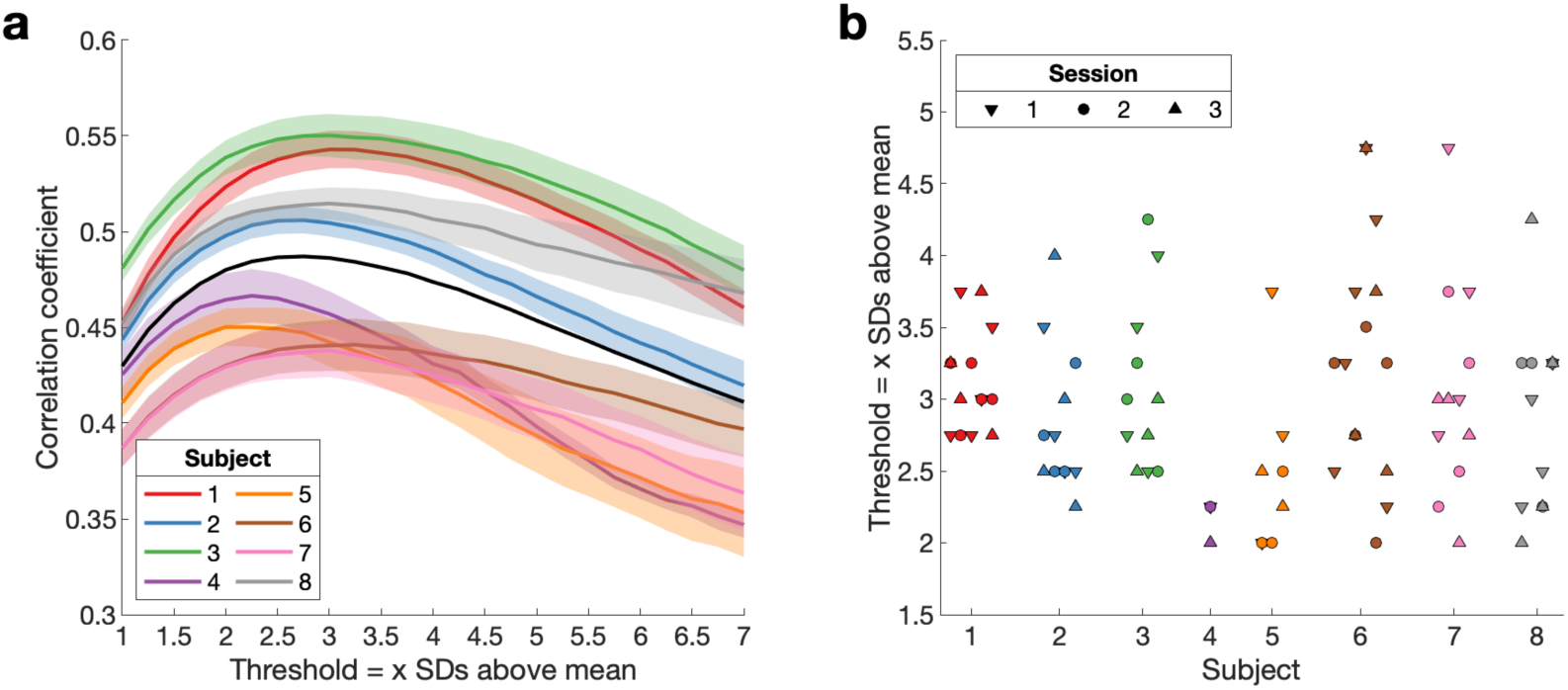
Empirical threshold to binarize beta bursts. To account for difference in signal-to-noise across sessions, days and subjects we obtained one threshold per session. (a) Mean correlation curves across days for each subject and sessions (+/- SEM) and across subjects (black line). (b) Empirical threshold for each session.

**Supplemental Fig. 3.**
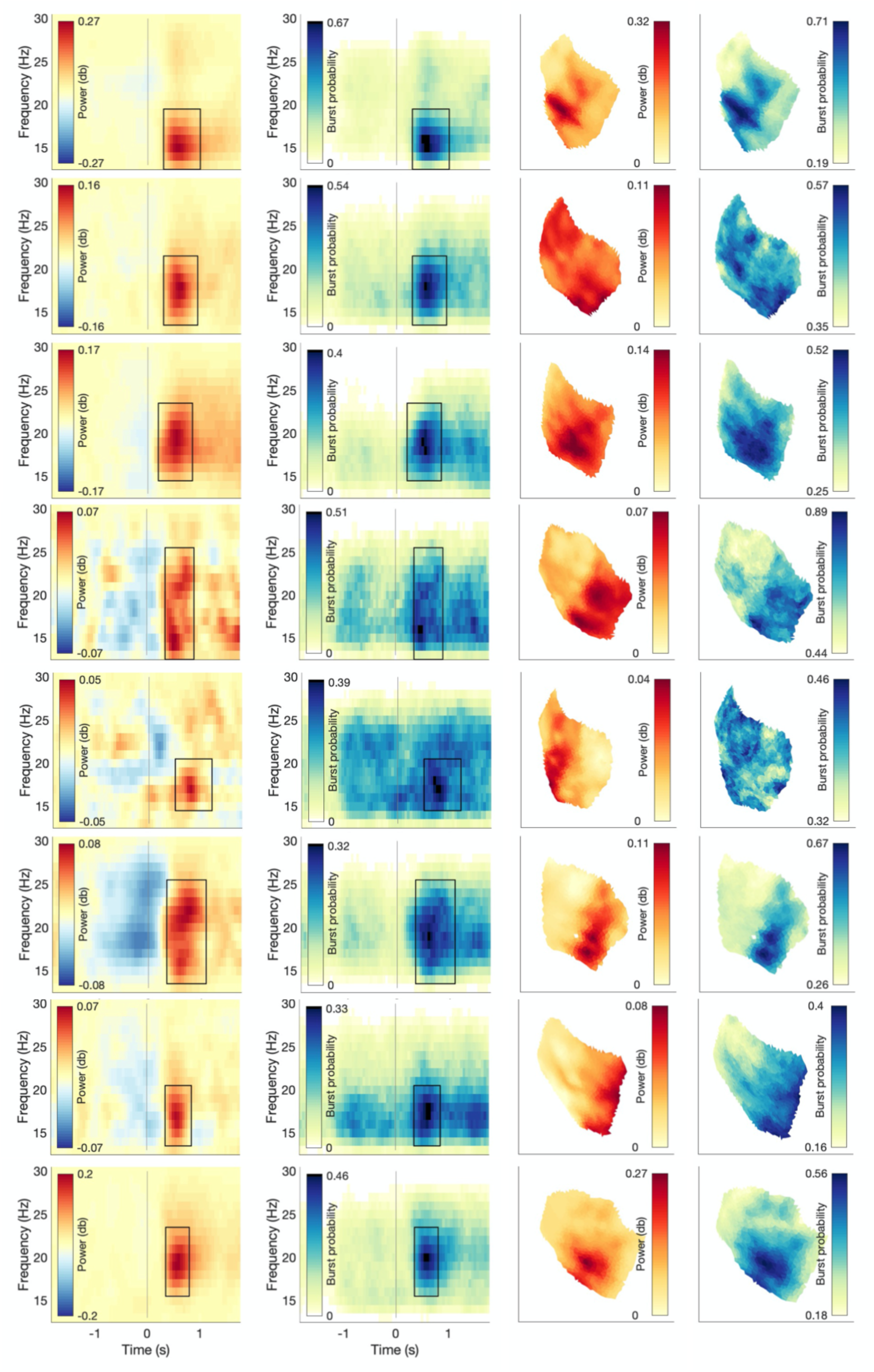
Beta power and burst probability are show for all three signal domains for each subject. (left) Conventional beta power and burst probability are shown as a function of time and frequency. To this end, data are averaged across the ROI. (right) Conventional beta power and burst probability as a function of space averaged in time and frequency (indicated by the rectangle in the time-frequency plot.

**Supplemental Fig. 4.**
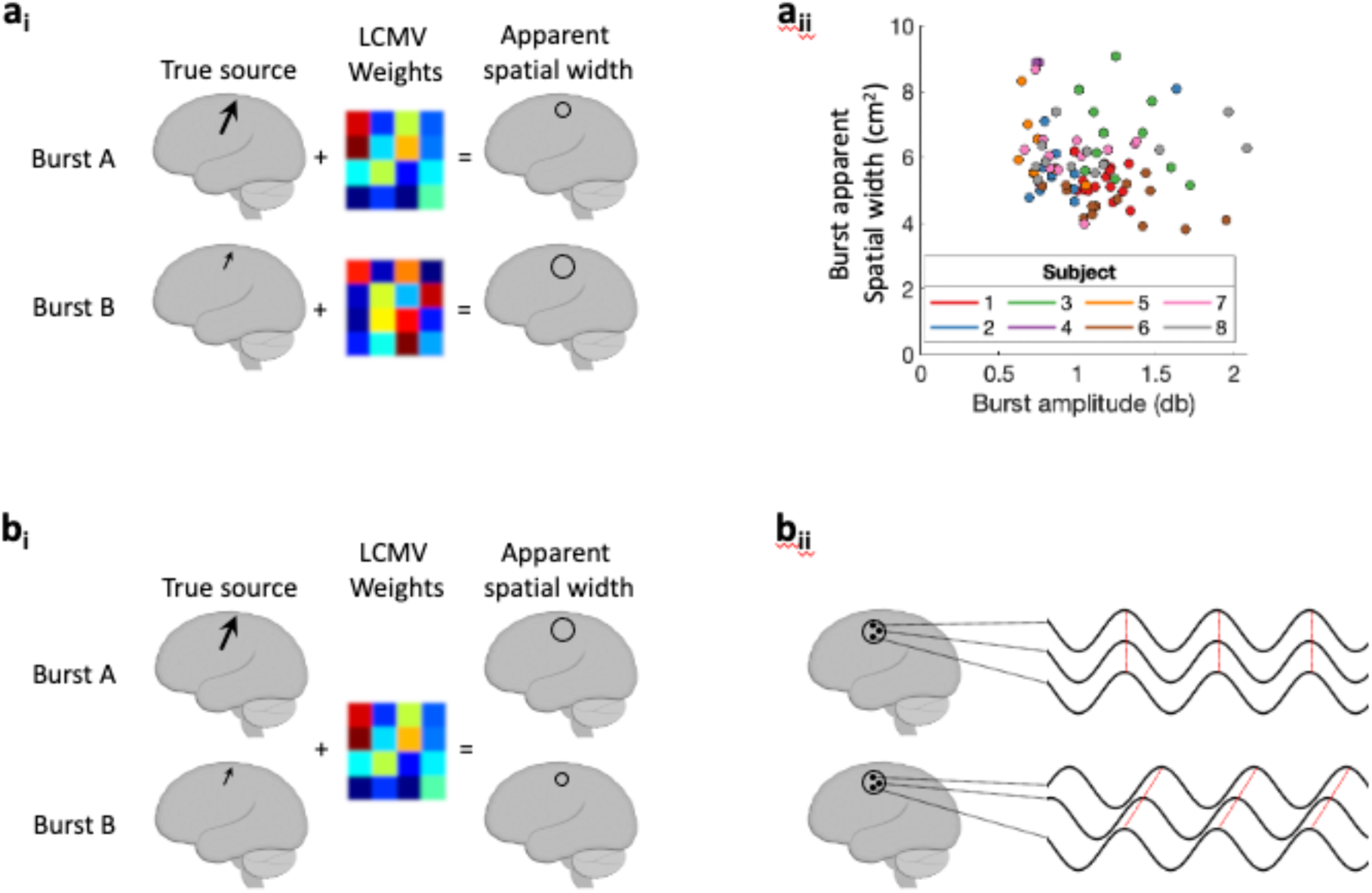
ai) Schematic illustration of how differences in SNR across sessions could theoretically explain variability in bursts’ apparent spatial width. Burst A (high amplitude) and burst B (low amplitude) each with distinct LCMV weights. If SNR across sessions explains the variability in bursts’ apparent spatial width, the apparent spatial width should be larger for small amplitude bursts. aii) Relationship between burst amplitude and burst apparent spatial width across sessions within and across subjects. bi) Schematic illustration of how differences in SNR within a session could theoretically explain variability in bursts’ apparent spatial width. Burst A (high amplitude) and burst B (low amplitude) with shared LCMV weights. If SNR within a session explains the variability in bursts’ apparent spatial width, the apparent spatial width should be larger for high amplitude bursts. bii) If bursts’ apparent spatial width is merely modulated by differences in SNR across bursts within a session, 1) a positive relationship between burst amplitude and burst apparent spatial width within sessions would be present, and 2) systematic phase differences across different spatial locations within each burst should be absent. Regarding the latter, if bursts’ apparent spatial width arises merely from amplitude scaling of a single source neural activity would show the same phase across different spatial locations of the burst (top). In turn, systematic phase lags across different spatial locations within the burst (bottom, Fig. 5) indicate that bursts’ apparent spatial width is unlikely to arise merely from amplitude scaling of a single source.

**Supplemental Fig. 5.**
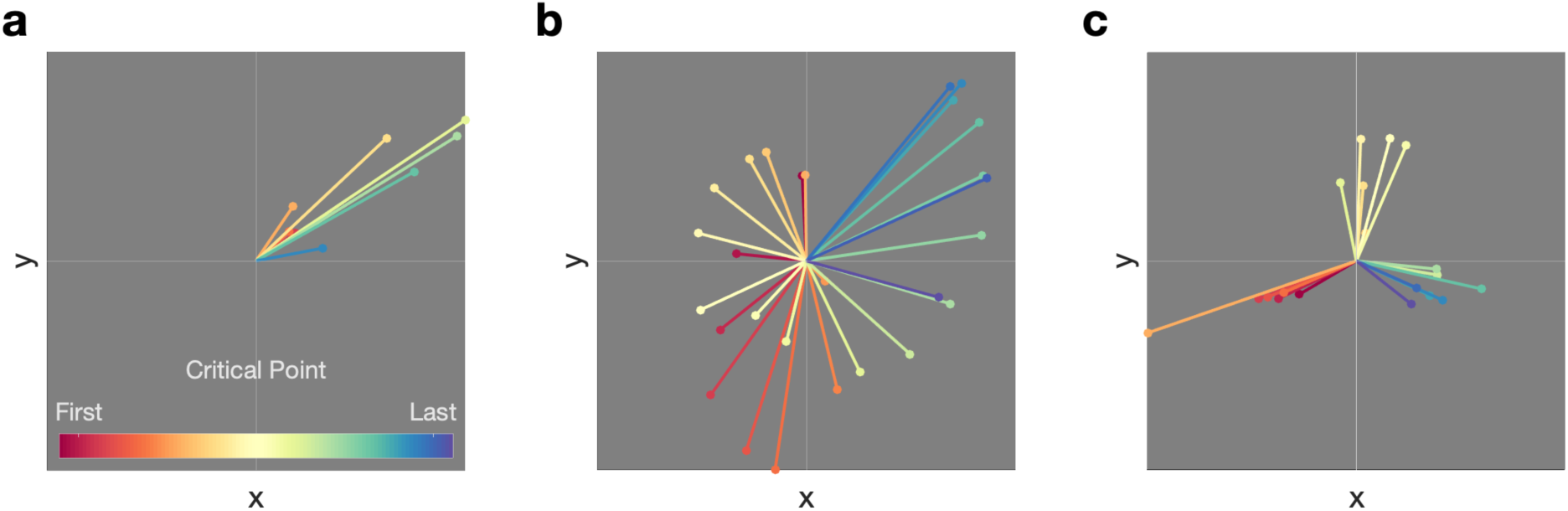
Examples of bursts with (a) one propagation direction and (b, c) complex propagation patterns.

**Supplemental Fig. 6.**
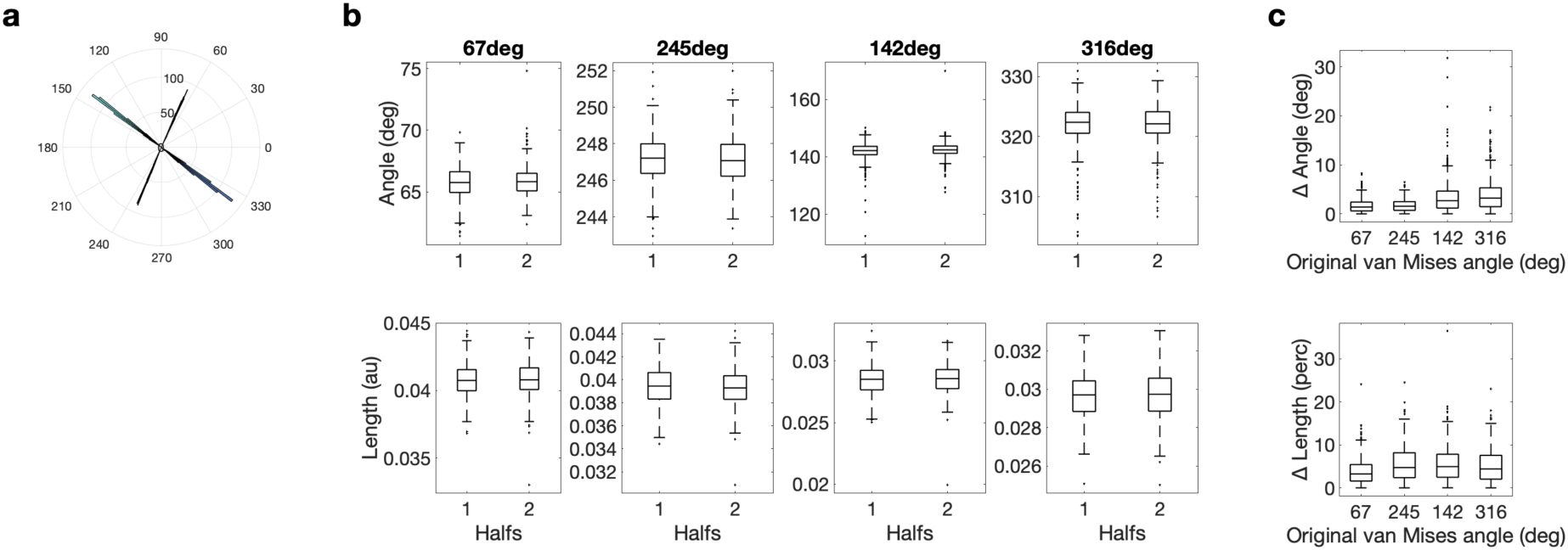
Length and angle are highly replicable for all four von Mises functions across halves of the data. (a) Histogram of the four van Mises functions across repetitions and halves. (b) Split-half reliability for angle (top) and length (bottom) for each van Mises function. (c) Difference between the two halves for all four van Mises functions. For angle the angular difference (top) and for length the percentage difference in length (bottom) is reported.

**Supplemental Fig. 7.**
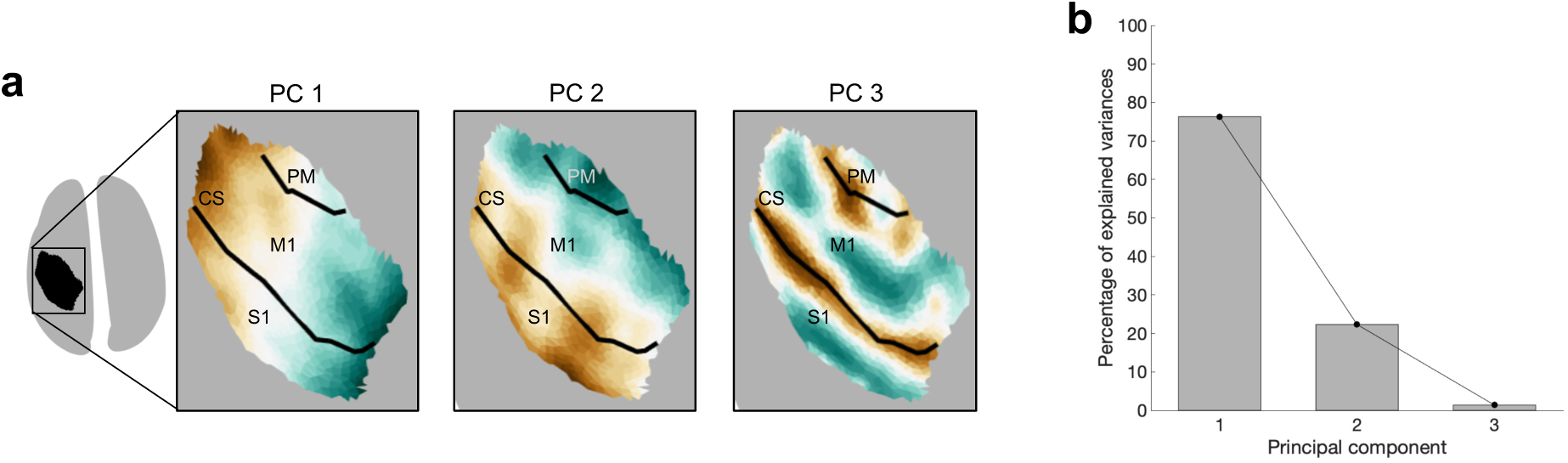
The spatial location of a burst can be summarized by the first two Principal Components (PCs) of the Cartesian coordinates of the centre of the burst. (a) For each PC the surface plot of the component structure is shown. CS, Central Sulcus. S1, Primary Sensory Cortex. M1, Primary Motor Cortex. PM, Premotor Cortex. (b) Variance explained by each principal component.

**Supplemental Fig. 8.**
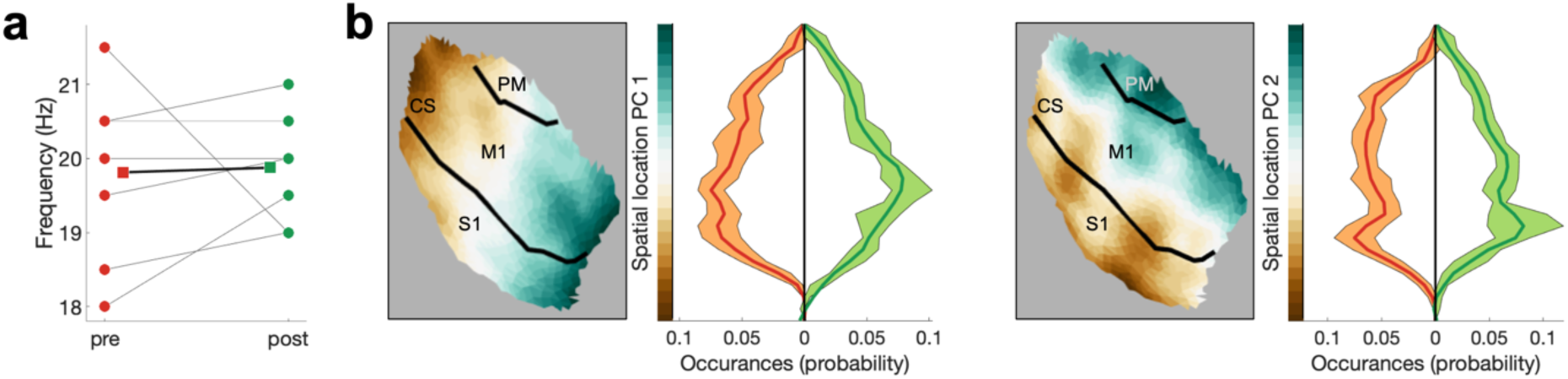
Frequency centre (a) and spatial location (b) and were not significantly different between pre-movement (red) and post-movement (green) bursts. CS, Central Sulcus. S1, Primary Sensory Cortex. M1, Primary Motor Cortex. PM, Premotor Cortex.

**Supplemental Fig. 9.**
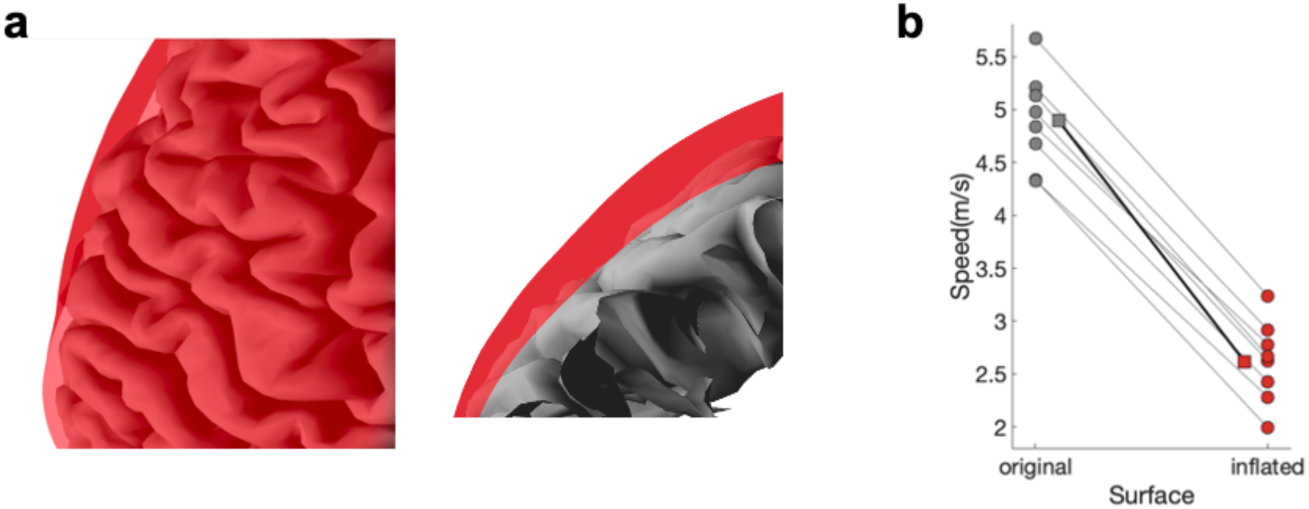
Propagation speed using the distance on the original or the inflated surface. (a) Overlay of the original (grey) and inflated surface (red). (b) Medians are shown for each subject (circles) and the mean across the subjects’ medians (square).

**Supplemental Fig. 10.**
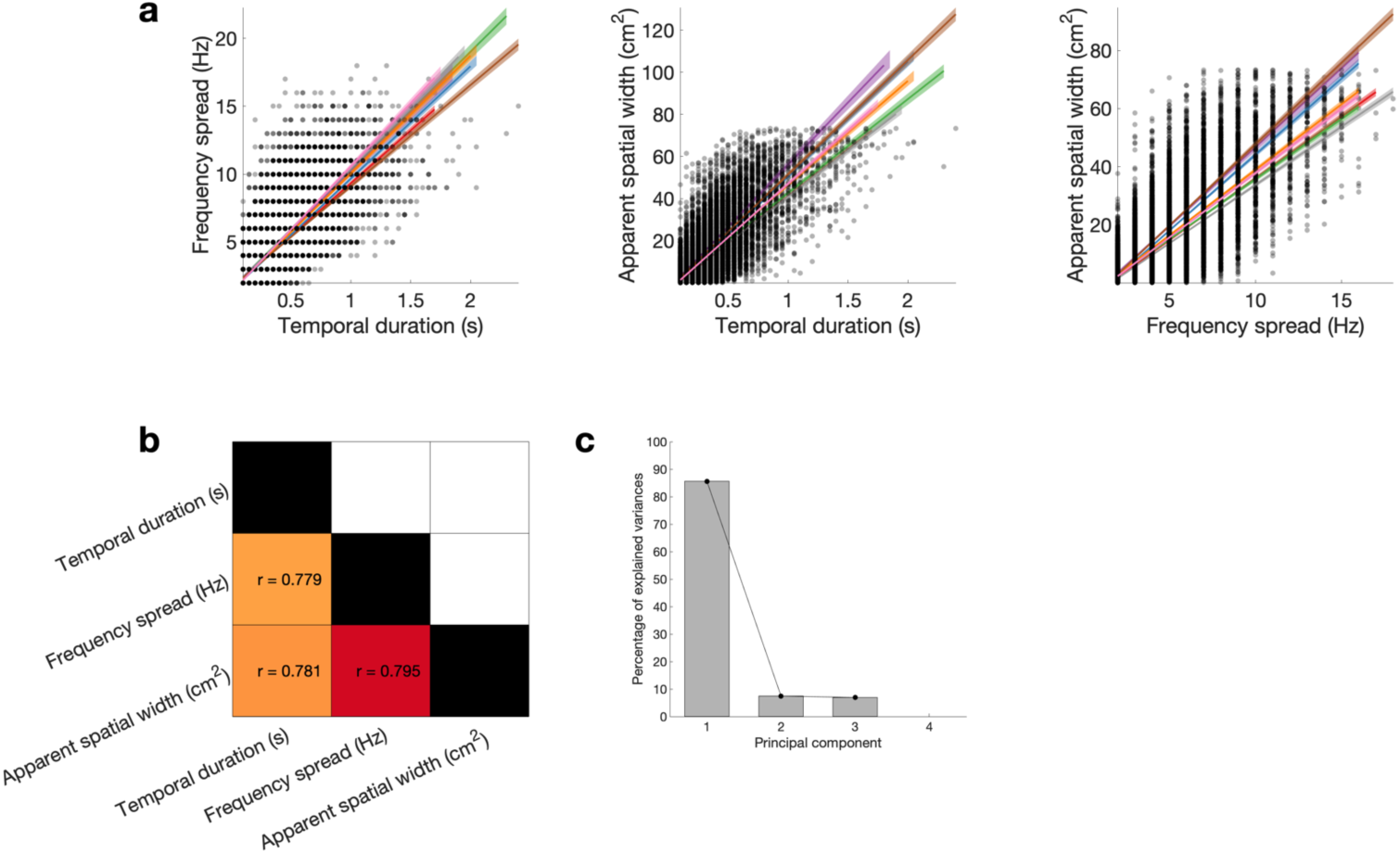
Reducing temporal duration, frequency spread and apparent spatial width to burst extend using Principal Component Analysis (PCA). (a) Temporal duration, frequency spread, and apparent spatial width are highly correlated across bursts. Coloured lines represent the least-squares fit for each subject and shaded areas indicate 95% confidence intervals. (b) Correlation matrix, whereby the Pearson correlation is averaged across subjects within each cell. (c) Variance explained by each principal component.

## References

Alexander, D. M., Ball, T., Schulze-Bonhage, A., & van Leeuwen, C. (2019). Large-scale cortical travelling waves predict localized future cortical signals. PLOS Computational Biology, 15(11), e1007316. https://doi.org/10.1371/journal.pcbi.1007316

Alexander, D. M., Nikolaev, A. R., Jurica, P., Zvyagintsev, M., Mathiak, K., & van Leeuwen, C. (2016). Global Neuromagnetic Cortical Fields Have Non-Zero Velocity. PLOS ONE, 11(3), e0148413. https://doi.org/10.1371/journal.pone.0148413

Bahramisharif, A., van Gerven, M. A. J., Aarnoutse, E. J., Mercier, M. R., Schwartz, T. H., Foxe, J. J., Ramsey, N. F., & Jensen, O. (2013). Propagating Neocortical Gamma Bursts Are Coordinated by Traveling Alpha Waves. Journal of Neuroscience, 33(48), 18849–18854. https://doi.org/10.1523/JNEUROSCI.2455-13.2013

Balasubramanian, K., Papadourakis, V., Liang, W., Takahashi, K., Best, M. D., Suminski, A. J., & Hatsopoulos, N. G. (2020). Propagating Motor Cortical Dynamics Facilitate Movement Initiation. Neuron, 106(3), 526–536.e4. https://doi.org/10.1016/j.neuron.2020.02.011

Best, M. D., Suminski, A. J., Takahashi, K., Brown, K. A., & Hatsopoulos, N. G. (2016). Spatio-Temporal Patterning in Primary Motor Cortex at Movement Onset. Cerebral Cortex, 27(2). https://doi.org/10.1093/cercor/bhv327

Bhattacharya, S., Brincat, S. L., Lundqvist, M., & Miller, E. K. (2022). Traveling waves in the prefrontal cortex during working memory. PLOS Computational Biology, 18(1), e1009827. https://doi.org/10.1371/journal.pcbi.1009827

Bonaiuto, J. J., Little, S., Neymotin, S. A., Jones, S. R., Barnes, G. R., & Bestmann, S. (2021). Laminar dynamics of high amplitude beta bursts in human motor cortex. NeuroImage, 242, 118479. https://doi.org/10.1016/j.neuroimage.2021.118479

Bonaiuto, J. J., Meyer, S. S., Little, S., Rossiter, H., Callaghan, M. F., Dick, F., Barnes, G. R., & Bestmann, S. (2018). Lamina-specific cortical dynamics in human visual and sensorimotor cortices. ELife, 7, e33977. https://doi.org/10.7554/eLife.33977

Brittain, J.-S., & Brown, P. (2014). Oscillations and the basal ganglia: Motor control and beyond. NeuroImage, 85, 637–647. https://doi.org/10.1016/j.neuroimage.2013.05.084

Burkitt, G. R., Silberstein, R. B., Cadusch, P. J., & Wood, A. W. (2000). Steady-state visual evoked potentials and travelling waves. Clinical Neurophysiology, 111(2), 246–258. https://doi.org/10.1016/S1388-2457(99)00194-7

Cagnan, H., Mallet, N., Moll, C. K. E., Gulberti, A., Holt, A. B., Westphal, M., Gerloff, C., Engel, A. K., Hamel, W., Magill, P. J., Brown, P., & Sharott, A. (2019). Temporal evolution of beta bursts in the parkinsonian cortical and basal ganglia network. Proceedings of the National Academy of Sciences, 116(32), 16095–16104. https://doi.org/10.1073/pnas.1819975116

Carey, D., Caprini, F., Allen, M., Lutti, A., Weiskopf, N., Rees, G., Callaghan, M. F., & Dick, F. (2018). Quantitative MRI provides markers of intra-, inter-regional, and age-related differences in young adult cortical microstructure. NeuroImage, 182, 429–440. https://doi.org/10.1016/j.neuroimage.2017.11.066

Carhart-Harris, R. L. (2018). The entropic brain—Revisited. Neuropharmacology, 142, 167–178. https://doi.org/10.1016/j.neuropharm.2018.03.010

Carhart-Harris, R. L., Leech, R., Hellyer, P. J., Shanahan, M., Feilding, A., Tagliazucchi, E., Chialvo, D. R., & Nutt, D. (2014). The entropic brain: A theory of conscious states informed by neuroimaging research with psychedelic drugs. Frontiers in Human Neuroscience, 8. https://doi.org/10.3389/fnhum.2014.00020

Cauller, L. J., Clancy, B., & Connors, B. W. (1998). Backward cortical projections to primary somatosensory cortex in rats extend long horizontal axons in layer I. The Journal of Comparative Neurology, 390(2), 297–310. https://doi.org/10.1002/(SICI)1096-9861(19980112)390:2<297::AID-CNE11>3.0.CO;2-V

Davis, Z. W., Benigno, G. B., Fletterman, C., Desbordes, T., Steward, C., Sejnowski, T. J., H. Reynolds, J., & Muller, L. (2021). Spontaneous traveling waves naturally emerge from horizontal fiber time delays and travel through locally asynchronous-irregular states. Nature Communications, 12(1), 6057. https://doi.org/10.1038/s41467-021-26175-1

Davis, Z. W., Muller, L., Martinez-Trujillo, J., Sejnowski, T., & Reynolds, J. H. (2020). Spontaneous travelling cortical waves gate perception in behaving primates. Nature, 587(7834), 432–436. https://doi.org/10.1038/s41586-020-2802-y

Deffains, M., Iskhakova, L., Katabi, S., Israel, Z., & Bergman, H. (2018). Longer β oscillatory episodes reliably identify pathological subthalamic activity in Parkinsonism: Longer STN β Episodes Are a PD Biomarker. Movement Disorders, 33(10), 1609–1618. https://doi.org/10.1002/mds.27418

Denker, M., Zehl, L., Kilavik, B. E., Diesmann, M., Brochier, T., Riehle, A., & Grün, S. (2018). LFP beta amplitude is linked to mesoscopic spatio-temporal phase patterns. Scientific Reports, 8(1), 5200. https://doi.org/10.1038/s41598-018-22990-7

Ding, Y., & Ermentrout, B. (2021). Traveling waves in non-local pulse-coupled networks. Journal of Mathematical Biology, 82(3), 18. https://doi.org/10.1007/s00285-021-01572-8

Dum, R. P. (2005). Frontal Lobe Inputs to the Digit Representations of the Motor Areas on the Lateral Surface of the Hemisphere. Journal of Neuroscience, 25(6), 1375–1386. https://doi.org/10.1523/JNEUROSCI.3902-04.2005

Engel, A. K., & Fries, P. (2010). Beta-band oscillations—Signalling the status quo? Current Opinion in Neurobiology, 20(2), 156–165. https://doi.org/10.1016/j.conb.2010.02.015

Enz, N., Ruddy, K. L., Rueda-Delgado, L. M., & Whelan, R. (2021). Volume of β-Bursts, But Not Their Rate, Predicts Successful Response Inhibition. The Journal of Neuroscience, 41(23), 5069–5079. https://doi.org/10.1523/JNEUROSCI.2231-20.2021

Ermentrout GB, K. (2001). Traveling Electrical Waves in Cortex: Insights from Phase Dynamics and Speculation on a Computational Role. 29, 33–44.

Feingold, J., Gibson, D. J., DePasquale, B., & Graybiel, A. M. (2015). Bursts of beta oscillation differentiate postperformance activity in the striatum and motor cortex of monkeys performing movement tasks. Proceedings of the National Academy of Sciences, 112(44), 13687–13692. https://doi.org/10.1073/pnas.1517629112

Fischl, B. (2012). FreeSurfer. NeuroImage, 62(2), 774–781. https://doi.org/10.1016/j.neuroimage.2012.01.021

Ghosh, S., & Gattera, R. (1995). A Comparison of the Ipsilateral Cortical Projections to the Dorsal and Ventral Subdivisions of the Macaque Premotor Cortex. Somatosensory & Motor Research, 12(3–4), 359–378. https://doi.org/10.3109/08990229509093668

Gilbertson, T., Lalo, E., Doyle, L., Di Lazzaro, V., Cioni, B., & Brown, P. (2005). Existing motor state is favored at the expense of new movement during 13-35 Hz oscillatory synchrony in the human corticospinal system. The Journal of Neuroscience: The Official Journal of the Society for Neuroscience, 25(34), 7771–7779. https://doi.org/10.1523/JNEUROSCI.1762-05.2005

Girard, P., Hupé, J. M., & Bullier, J. (2001). Feedforward and feedback connections between areas V1 and V2 of the monkey have similar rapid conduction velocities. Journal of Neurophysiology, 85(3), 1328–1331. https://doi.org/10.1152/jn.2001.85.3.1328

Heideman, S. G., Quinn, A. J., Woolrich, M. W., van Ede, F., & Nobre, A. C. (2020). Dissecting beta-state changes during timed movement preparation in Parkinson’s disease. Progress in Neurobiology, 184, 101731. https://doi.org/10.1016/j.pneurobio.2019.101731

Heitmann, S., Rule, M., Truccolo, W., & Ermentrout, B. (2017). Optogenetic Stimulation Shifts the Excitability of Cerebral Cortex from Type I to Type II: Oscillation Onset and Wave Propagation. PLOS Computational Biology, 13(1), e1005349. https://doi.org/10.1371/journal.pcbi.1005349

Hughes, J. R. (1995). The Phenomenon of Travelling Waves: A Review. Clinical Electroencephalography, 26(1), 1–6. https://doi.org/10.1177/155005949502600103

Hurtado, J. M., Rubchinsky, L. L., & Sigvardt, K. A. (2004). Statistical Method for Detection of Phase-Locking Episodes in Neural Oscillations. Journal of Neurophysiology, 91(4), 1883–1898. https://doi.org/10.1152/jn.00853.2003

Hyvarinen, A. (1999). Fast and robust fixed-point algorithms for independent component analysis. IEEE Transactions on Neural Networks, 10(3), 626–634. https://doi.org/10.1109/72.761722

Jones, S. R. (2016). When brain rhythms aren’t ‘rhythmic’: Implication for their mechanisms and meaning. Current Opinion in Neurobiology, 40, 72–80. https://doi.org/10.1016/j.conb.2016.06.010

Joundi, R. A., Jenkinson, N., Brittain, J.-S., Aziz, T. Z., & Brown, P. (2012). Driving Oscillatory Activity in the Human Cortex Enhances Motor Performance. Current Biology, 22(5), 403–407. https://doi.org/10.1016/j.cub.2012.01.024

Kaufman, M. T., Churchland, M. M., Ryu, S. I., & Shenoy, K. V. (2014). Cortical activity in the null space: Permitting preparation without movement. Nature Neuroscience, 17(3), 440–448. https://doi.org/10.1038/nn.3643

Khanna, P., & Carmena, J. M. (2017). Beta band oscillations in motor cortex reflect neural population signals that delay movement onset. ELife, 6, e24573. https://doi.org/10.7554/eLife.24573

Kurata, K. (1991). Corticocortical inputs to the dorsal and ventral aspects of the premotor cortex of macaque monkeys. Neuroscience Research, 12(1), 263–280. https://doi.org/10.1016/0168-0102(91)90116-G

Landler, L., Ruxton, G. D., & Malkemper, E. P. (2021). Advice on comparing two independent samples of circular data in biology. Scientific Reports, 11(1), 20337. https://doi.org/10.1038/s41598-021-99299-5

Law, R. G., Pugliese, S., Shin, H., Sliva, D. D., Lee, S., Neymotin, S., Moore, C., & Jones, S. R. (2022). Thalamocortical Mechanisms Regulating the Relationship between Transient Beta Events and Human Tactile Perception. Cerebral Cortex, 32(4), 668–688. https://doi.org/10.1093/cercor/bhab221

Little, S., Bonaiuto, J., Barnes, G., & Bestmann, S. (2019). Human motor cortical beta bursts relate to movement planning and response errors. PLOS Biology, 17(10), e3000479. https://doi.org/10.1371/journal.pbio.3000479

Luppino, G., Matelli, M., Camarda, R., & Rizzolatti, G. (1993). Corticocortical connections of area F3 (SMA-proper) and area F6 (pre-SMA) in the macaque monkey. The Journal of Comparative Neurology, 338(1), 114–140. https://doi.org/10.1002/cne.903380109

Luppino, G., & Rizzolatti, G. (2000). The Organization of the Frontal Motor Cortex. Physiology, 15(5), 219–224. https://doi.org/10.1152/physiologyonline.2000.15.5.219

Meyer, S. S., Bonaiuto, J., Lim, M., Rossiter, H., Waters, S., Bradbury, D., Bestmann, S., Brookes, M., Callaghan, M. F., Weiskopf, N., & Barnes, G. R. (2017). Flexible head-casts for high spatial precision MEG. Journal of Neuroscience Methods, 276, 38–45. https://doi.org/10.1016/j.jneumeth.2016.11.009

Muakkassa, K. F., & Strick, P. L. (1979). Frontal lobe inputs to primate motor cortex: Evidence for four somatotopically organized ‘premotor’ areas. Brain Research, 177(1), 176–182. https://doi.org/10.1016/0006-8993(79)90928-4

Muller, L., Chavane, F., Reynolds, J., & Sejnowski, T. J. (2018). Cortical travelling waves: Mechanisms and computational principles. Nature Reviews Neuroscience, 19(5), 255–268. https://doi.org/10.1038/nrn.2018.20

Neymotin, S. A., Daniels, D. S., Caldwell, B., McDougal, R. A., Carnevale, N. T., Jas, M., Moore, C. I., Hines, M. L., Hämäläinen, M., & Jones, S. R. (2020). Human Neocortical Neurosolver (HNN), a new software tool for interpreting the cellular and network origin of human MEG/EEG data. ELife, 9, e51214. https://doi.org/10.7554/eLife.51214

Penfield, W., & Boldrey, E. (1937). Somatic motor and sensory representation in the cerebral cortex of man studied by electrical stimulation. Brain, 60(4), 389–443. https://doi.org/10.1093/brain/60.4.389

Pogosyan, A., Gaynor, L. D., Eusebio, A., & Brown, P. (2009). Boosting Cortical Activity at Beta-Band Frequencies Slows Movement in Humans. Current Biology, 19(19), 1637–1641. https://doi.org/10.1016/j.cub.2009.07.074

Prechtl, J. C., Bullock, T. H., & Kleinfeld, D. (2000). Direct evidence for local oscillatory current sources and intracortical phase gradients in turtle visual cortex. Proceedings of the National Academy of Sciences, 97(2), 877–882. https://doi.org/10.1073/pnas.97.2.877

Quinn, A. J., van Ede, F., Brookes, M. J., Heideman, S. G., Nowak, M., Seedat, Z. A., Vidaurre, D., Zich, C., Nobre, A. C., & Woolrich, M. W. (2019). Unpacking Transient Event Dynamics in Electrophysiological Power Spectra. Brain Topography, 32(6), 1020–1034. https://doi.org/10.1007/s10548-019-00745-5

Riehle, A., Wirtssohn, S., Grün, S., & Brochier, T. (2013). Mapping the spatio-temporal structure of motor cortical LFP and spiking activities during reach-to-grasp movements. Frontiers in Neural Circuits, 7. https://doi.org/10.3389/fncir.2013.00048

Roberts, J. A., Gollo, L. L., Abeysuriya, R. G., Roberts, G., Mitchell, P. B., Woolrich, M. W., & Breakspear, M. (2019). Metastable brain waves. Nature Communications, 10(1), 1056. https://doi.org/10.1038/s41467-019-08999-0

Rosner, B. (1983). Percentage Points for a Generalized ESD Many-Outlier Procedure. Technometrics, 25(2), 9.

Rubino, D., Robbins, K. A., & Hatsopoulos, N. G. (2006). Propagating waves mediate information transfer in the motor cortex. Nature Neuroscience, 9(12), 1549–1557. https://doi.org/10.1038/nn1802

Rule, M. E., Vargas-Irwin, C., Donoghue, J. P., & Truccolo, W. (2018). Phase reorganization leads to transient β-LFP spatial wave patterns in motor cortex during steady-state movement preparation. Journal of Neurophysiology, 119(6), 2212–2228. https://doi.org/10.1152/jn.00525.2017

Seedat, Z. A., Quinn, A. J., Vidaurre, D., Liuzzi, L., Gascoyne, L. E., Hunt, B. A. E., O’Neill, G. C., Pakenham, D. O., Mullinger, K. J., Morris, P. G., Woolrich, M. W., & Brookes, M. J. (2020). The role of transient spectral ‘bursts’ in functional connectivity: A magnetoencephalography study. NeuroImage, 209, 116537. https://doi.org/10.1016/j.neuroimage.2020.116537

Sherman, M. A., Lee, S., Law, R., Haegens, S., Thorn, C. A., Hämäläinen, M. S., Moore, C. I., & Jones, S. R. (2016). Neural mechanisms of transient neocortical beta rhythms: Converging evidence from humans, computational modeling, monkeys, and mice. Proceedings of the National Academy of Sciences, 113(33), E4885–E4894. https://doi.org/10.1073/pnas.1604135113

Shin, H., Law, R., Tsutsui, S., Moore, C. I., & Jones, S. R. (2017). The rate of transient beta frequency events predicts behavior across tasks and species. ELife, 6, e29086. https://doi.org/10.7554/eLife.29086

Sporn, S., Hein, T., & Herrojo Ruiz, M. (2020). Alterations in the amplitude and burst rate of beta oscillations impair reward-dependent motor learning in anxiety. ELife, 9, e50654. https://doi.org/10.7554/eLife.50654

Sreekumar, V., Wittig, J. H., Chapeton, J. I., Inati, S. K., & Zaghloul, K. A. (2020). *Low frequency traveling waves in the human cortex coordinate neural activity across spatial scales* [Preprint]. Neuroscience. https://doi.org/10.1101/2020.03.04.977173

Stolk, A., Brinkman, L., Vansteensel, M. J., Aarnoutse, E., Leijten, F. S., Dijkerman, C. H., Knight, R. T., de Lange, F. P., & Toni, I. (2019). Electrocorticographic dissociation of alpha and beta rhythmic activity in the human sensorimotor system. ELife, 8, e48065. https://doi.org/10.7554/eLife.48065

Swadlow, & Waxman. (2012). Axonal conduction delays. Scholarpedia, 7(1451).

Takahashi, K., Kim, S., Coleman, T. P., Brown, K. A., Suminski, A. J., Best, M. D., & Hatsopoulos, N. G. (2015). Large-scale spatiotemporal spike patterning consistent with wave propagation in motor cortex. Nature Communications, 6(1), 7169. https://doi.org/10.1038/ncomms8169

Takahashi, K., Saleh, M., Penn, R. D., & Hatsopoulos, N. G. (2011). Propagating Waves in Human Motor Cortex. Frontiers Human Neuroscience, 5. https://doi.org/10.3389/fnhum.2011.00040

Tinkhauser, G., Pogosyan, A., Little, S., Beudel, M., Herz, D. M., Tan, H., & Brown, P. (2017). The modulatory effect of adaptive deep brain stimulation on beta bursts in Parkinson’s disease. Brain, 140(4), 1053–1067. https://doi.org/10.1093/brain/awx010

Tinkhauser, G., Pogosyan, A., Tan, H., Herz, D. M., Kühn, A. A., & Brown, P. (2017). Beta burst dynamics in Parkinson’s disease OFF and ON dopaminergic medication. Brain, 140(11), 2968–2981. https://doi.org/10.1093/brain/awx252

Troebinger, L., López, J. D., Lutti, A., Bradbury, D., Bestmann, S., & Barnes, G. (2014). High precision anatomy for MEG. NeuroImage, 86, 583–591. https://doi.org/10.1016/j.neuroimage.2013.07.065

Van Veen, B. D., & Buckley, K. M. (1988). Beamforming: A versatile approach to spatial filtering. IEEE ASSP Magazine, 5(2), 4–24. https://doi.org/10.1109/53.665

Vidaurre, D., Hunt, L. T., Quinn, A. J., Hunt, B. A. E., Brookes, M. J., Nobre, A. C., & Woolrich, M. W. (2018). Spontaneous cortical activity transiently organises into frequency specific phase-coupling networks. Nature Communications, 9(1), 2987. https://doi.org/10.1038/s41467-018-05316-z

Wessel, J. R. (2020). β-Bursts Reveal the Trial-to-Trial Dynamics of Movement Initiation and Cancellation. The Journal of Neuroscience, 40(2), 411–423. https://doi.org/10.1523/JNEUROSCI.1887-19.2019

Woolrich, M., Hunt, L., Groves, A., & Barnes, G. (2011). MEG beamforming using Bayesian PCA for adaptive data covariance matrix regularization. NeuroImage, 57(4), 1466–1479. https://doi.org/10.1016/j.neuroimage.2011.04.041

Zich, C., Quinn, A. J., Mardell, L. C., Ward, N. S., & Bestmann, S. (2020). Dissecting Transient Burst Events. Trends in Cognitive Sciences, 24(10), 784–788. https://doi.org/10.1016/j.tics.2020.07.004

Zich, C., Woolrich, M. W., Becker, R., Vidaurre, D., Scholl, J., Hinson, E. L., Josephs, L., Braeutigam, S., Quinn, A. J., & Stagg, C. J. (2018). Motor learning shapes temporal activity in human sensorimotor cortex [Preprint]. Neuroscience. https://doi.org/10.1101/345421

## References

Vidaurre, D., Quinn, A. J., Baker, A. P., Dupret, D., Tejero-Cantero, A., & Woolrich, M. W. (2016). Spectrally resolved fast transient brain states in electrophysiological data. NeuroImage, 126, 81–95. https://doi.org/10.1016/j.neuroimage.2015.11.047

